# Overexpression of barley heat stress transcription factor HvHsfA6a provide thermotolerance by thermopriming

**DOI:** 10.1101/2024.03.06.583688

**Authors:** Reeku Chaudhary, Suchi Baliyan, Parul Sirohi, Sweta Singh, Sumit Kumar Mishra, Mohan Singh Rajkumar, Shashank Sagar Saini, Hugo Germain, Debabrata Sircar, Harsh Chauhan

**Author notes:** Email of authors: Reeku Chaudhary, Suchi Baliyan, Parul Sirohi, Sweta Singh, Sumit Kumar Mishra, Mohan Singh Rajkumar, Shashank Sagar Saini, Hugo Germain, Debabrata Sircar, Harsh Chauhan.

## Abstract

Adverse impacts of climate change, including high temperature on cereal crop production, have been evidenced globally. In plants, heat shock factors (HSFs) are crucial components of heat stress associated rescue mechanisms and are also required for normal biological processes. Here, we functionally characterized a highly heat stress responsive *HvHSFA6a* in barley by developing constitutively overexpressing transgenic lines. These transgenic lines showed heat tolerant phenotype via improved photosynthesis, antioxidants and upregulation of HSPs and metabolites involved in stress amelioration and keeping thermomemory as compared to wild type plants. Global transcriptomics and ChIP sequencing revealed that HvHSFA6a orchestrates the expression of several genes through direct binding with other HSFs containing consensus HSE in their promoter regions. A GC-MS based metabolomics analysis also revealed the alterations in key metabolic processes such as carbohydrate metabolism, citric acid cycle, amino acids and secondary metabolism. Higher accumulation of key metabolites such as sucrose, galactinol, shikimate and ascorbate has been observed under both control and heat stress in transgenic lines as compared to wild type plants. Taken together, the results suggest that overexpression of *HvHsfA6a* prime the plants for heat stress conditions by alteration in gene expression and metabolic status.

**Highlight:** Priming is a mechanism by which plants respond to various abiotic and biotic stresses. Through multi omics approach we found that barley HsfA6a provide thermotolernce in transgenic plants through priming effect on transcriptome and metabolome.

## Introduction

Global climate change and rising temperature have direct adverse impacts on both quality of produce and yield of crop plants including cereals which are staple food for world population. The drop in the yield of cereals due to high temperature stress, leads to surges in food prices, thereby making global food security a challenge (FAO, 2018). Such challenges justify more efforts to generate high temperature resilient crops for which a detailed knowledge of pathways and genes associated with heat stress tolerance in plants is imperative.

Heat stress has effects across all events of plant life cycle i.e. from seed germination to seed maturation (Porter and Moot, 1998). Plants have remarkable ability to survive in stressful conditions including high temperature. In response to heat stress they evoke an interconnected signalling cascade and initiate heat shock response (HSR) to provide thermotolerance in plants. It has been shown that the phase transition of lipid bilayer triggers the heat stress-associated signalling pathways through calcium ions (Finka et al., 2012). These calcium ions are also known to govern the activity of heat shock factors (HSFs), which are key components of heat stress signalling cascade (Li et al., 2004). HSFs confer thermotolerance to plants through alterations in the transcriptional program of a set of heat shock responsive genes and control the expression of approximately 240 genes as shown in model plants such as Arabidopsis and tomato (Scharf et al., 2012; Liu and Charng, 2013; Fragkostefanakis et al., 2015).

In plants, HSF family is highly diversified based on their molecular structure and functional characteristics (von Koskull-Doring et al., 2007). HSFs are classified into three classes i.e. A, B and C on the basis of their modular structure and motif composition (Nover et al., 1996; Nover et al., 2001; Baniwal et al., 2004). Accordingly, Nover et al. (2001), classified Arabidopsis HSFs class A into 9 groups (A1-A9), and their number ranged from 21 in Arabidopsis, 25 in rice, 22 in barley, to 56 in wheat which is currently highest among plants (Nover et al., 2001; Chauhan et al., 2011; Reddy et al., 2014; Xue et al., 2014). Notably, HSFs possess unique functions as revealed by several expression and functional studies (von Koskull-Doring et al., 2007; Scharf et al., 2012; Wang et al., 2017). For example, HsfA1a has been reported as a master regulator of heat shock response in tomato, however, in Arabidopsis this role is shared by four isoforms of HsfA1 showing functional redundancy (Mishra et al., 2002; Liu et al., 2011). In tomato HsfA2 and HsfA1 interact synergistically to regulate the expression of their downstream genes (Chan-Schaminet et al., 2009).

Genetic studies including overexpression and mutant analysis of class A2 *HSFs* such as *AtHsfA2, OsHsfA2a, OsHsfA2e*, *TaHsfA2d*, *AtHsfA2, FaHsfA2c* from Arabidopsis, rice, wheat, lily and tall fescue respectively, revealed their role in conferring tolerance against heat and other abiotic stresses such as salt, osmotic, anoxia and oxidative stress (Schramm et al., 2006; Yokotani et al., 2007; Nishizawa-Yokoi et al., 2009; Zhang et al., 2009; Ogawa et al., 2007; Nishizawa et al., 2006; Banti et al., 2010; Chauhan et al., 2013).Furthermore, transcriptome analysis of transgenic *Arabidopsis* plants revealed that downstream genes of these HSFs included *heat shock protein* genes, *ascorbate peroxidase*, *galactinol synthase,* and other thermomemory related genes, to name a few (e.g. *HSPs*, *Apx* etc.). HsfA2 positively regulates the expression of these memory genes via chromatin modification (H3K4 methylations) and leads to prolonged activation of these genes (Liu et al., 2018). To execute thermomemory these Hsfs work independently and could form hetromeric complexes such as HsfA2/HsfA3 to induce thermomemory in Arabidopsis plants (Lamke et al., 2016, Friedrich et al., 2023). Apart from heat tolerance, other HSF of class A, such as *AtHsfA6a* conferred salinity and drought tolerance to transgenic Arabidopsis plants by activating ABA-mediated pathways, while being susceptible to heat (Hwang et al., 2019). In addition, overexpression study of *TaHsfA6f* under constitutive and drought inducible promoter in Arabidopsis and wheat, respectively, suggested that this HSF could bind to genes such as *Rubisco activase* (large isoform), Golgi anti-apoptotic protein (GAAP) alongwith HSPs and other HSR genes to confer tolerance towards heat, drought and salinity (Xue et al., 2015). HSFs bound to the promoter regions of heat shock elements, CArG, G-box, E-box containing motifs of stress responsive genes, to regulate primarily the heat shock response, as evidenced by genome wide ChIP-seq analysis, and mediate heat, salinity and drought tolerance in Arabidopsis plants (Zang et al., 2019, Albihlal et al., 2018).

Besides transcriptomic alterations HSFs also lead to metabolomic changes in plants during stressed environment. Song et al., (2016) found increased level of galactinol and its derivatives raffinose and stachyose in Arabidopsis plants during heat and oxidative stress. Whereas, metabolome of heat primed plants as compared to nonprimed plants showed increased synthesis of compounds involved in lipid and carbohydrate metabolism, tocopherols, and other thermomemory associated metabolites such as galactinol and raffinose (Serrano et al., 2019). These metabolic alteration aids plants to establish enhanced HS tolerance under repetitive stress.

However, despite extensive studies of HSFs in model plant Arabidopsis, till date such studies to understand the effect of HSFs on overall transcriptional regulation coupled with underlying metabolic dynamics are elusive in cereal crop plants. Therefore, we characterized the HSF of Class A namely *HvHsfA6a* in barley, which is considered as a model crop for *Triticeae* tribe for functional genomics studies (Saisho and Takeda, 2011) being highly susceptible to heat stress. In the present study, we constructed *HvHsfA6a* overexpressing transgenic lines in barley, using a constitutive *Zea mays Ubiquitin promoter* (*ZmUbi*). We further coupled transcriptome wide survey with chromatin immunoprecipitation sequencing assay to identify its genetic targets and to understand its mode of genetic regulation during heat stress. GC-MS based metabolomics analysis was performed to gain insights in metabolic alterations during HS in transgenics. Our results combining RNA-Seq and ChIP-Seq suggested that *HvHsfA6a* leads to the transcriptional reprogramming of genes associated with HSPs, photosynthetic pathways including both light and dark reactions, sugar metabolism alongwith normal growth and development associated genes. GC-MS based analysis furthermore confirmed these transcriptomic alterations at the metabolic level. The transgenic plants demonstrated no adverse phenotypic alteration and showed excellent performance under elevated temperature as compared to wild type plants. We propose that *HvHsfA6a* gene could be a potential candidate gene in providing thermotolerance in case of the cereal crop barley.

## RESULTS

### HvHsfA6a is heat-inducible, possesses transactivation property, and is nuclear localized

*HvHsfA6a* (AK358460) gene encodes 372 aa protein with all the characteristic features of class A Hsf members such as a DNA binding domain, an oligomerization domain, a nuclear localization signal, a transactivation domain (AHA1) and a nuclear export signal (Fig. 1A). At the transcript level, we noticed activation of this gene in the shoot (61-fold) and root (15-fold) tissues upon heat stress treatment (Fig. 1B). To check if HvHsfA6a possess transactivation property, like other Hsf members, we used a yeast one hybrid system. The yeast cells transformed with the ORF of *HvHsfA6a*, grew well on the medium of SD/−Trp, and the colonies appeared blue on the SD/−Trp medium supplied with X-α-Gal, indicating that HvHsfA6a is a transcriptional activator (Fig. 1C). HSFs have been reported to be both cytosolic and nuclear localized and shuttle between cytosol and nucleus upon exposure to heat stress (Scharf et al. (2012).

**Fig. 1.**
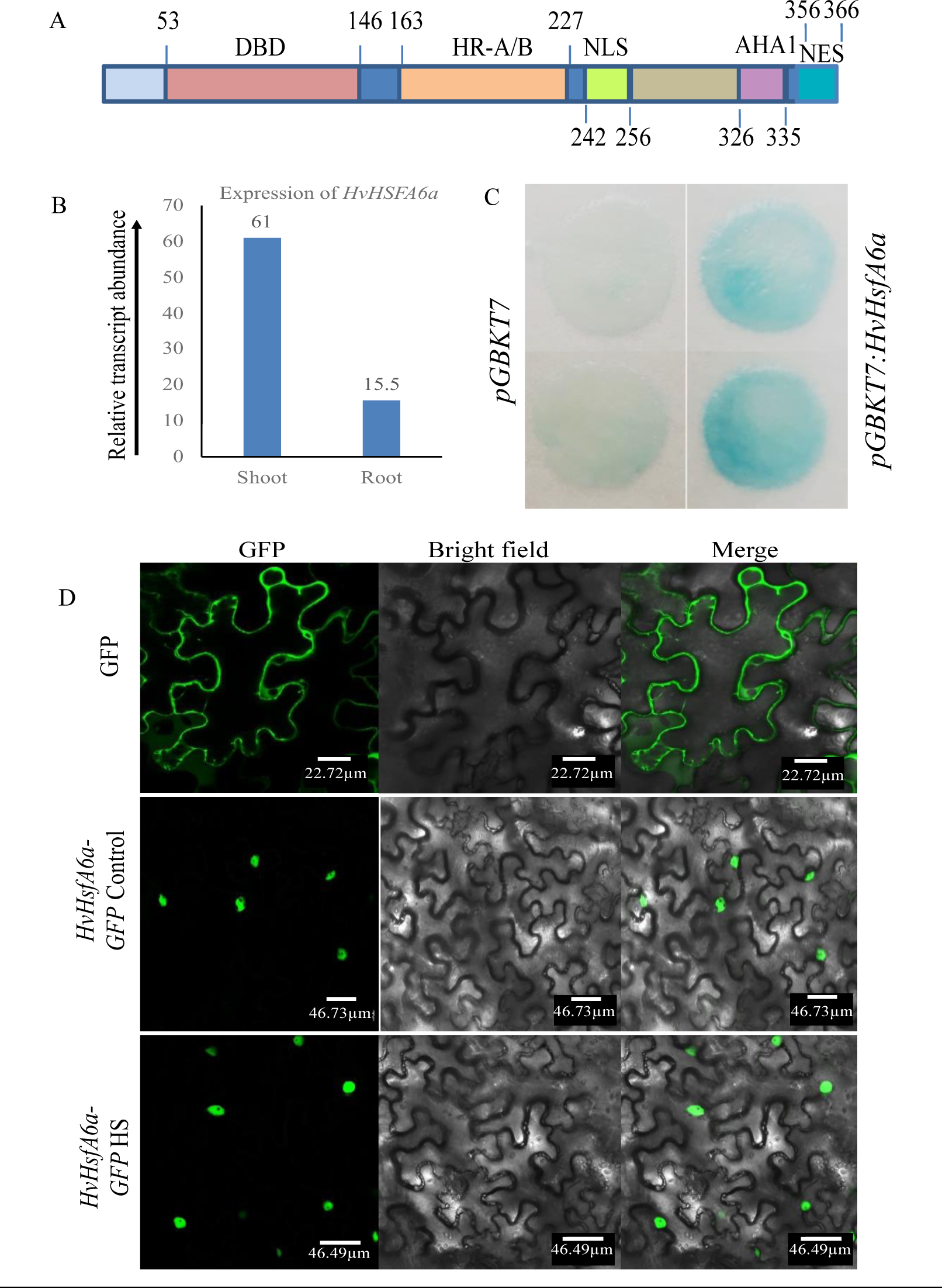
**HvHsfA6a is heat-inducible, transcriptional activator and nuclear localized** A) The modular structure of HvHSFA6a shows all the characteristic features of a typical class A Hsf, B) B) qRT-PCR-based expression profile of *HvHSFA6a* in both shoot and root tissues of barley after 15_o_ min of heat stress at 37 C. C) Transactivation potential of HvHsfA6a in GAL4BD filter lift assay. D) C-terminal GFP fused construct HvHSFA6a:GFP was found nuclear localized under controlled and during heat stress conditions.

Therefore, to assess the subcellular localization of HvHsfA6a during both normal and heat-stressed (HS) conditions, a C-terminal GFP fused construct with *HvHsfA6a* was prepared and transiently expressed in *Nicotiana benthamiana* leaves. HvHsfA6a:GFP fusion protein was found to be localized in the nucleus regardless of the presence of heat stress (Fig. 1D).

### Development of transgenic barley plants overexpressing *HvHsfA6a*

We next generated *HvHsfA6a* overexpression transgenic plants to elucidate its function in barley (Hensel et al., 2009). For this purpose, *HvHsfA6a* was cloned under *Zea mays ubiquitin promoter* (*ZmUbi1*) in the modified binary vector *pANIC6B* (Mann et al., 2012). We obtained a total of 50 transgenic lines named as *ZmUbi:HvHsfA6aRC* with varied levels of overexpression, as revealed by 5-20 -fold upregulation of *HvHsfA6a* in the qRT-PCR analysis. Further, immunoblot analysis was performed by using antiHvHSFA6a antibodies in selected transgenic lines exhibiting the highest level of overexpression at transcript levels to assess corresponding protein accumulation. As expected, the HvHSFA6a OX transgenic plants accumulated this protein constitutively, even under unstress conditions. In contrast, a heat-stressed inducible accumulation of this protein was witnessed in WT seedlings (Supplementary Fig. 1A). Based on their initial confirmation for HSFA6a overexpression, we selected RC8, RC16, RC19, and RC20 lines for morphological, transcriptome, biochemical and metabolomic analysis.

### *ZmUbi:HvHsfA6a* OX plants show improved tolerance to heat stress

To examine the heat stress tolerance potential of RC8, RC16, RC19 and RC20 lines, wild-type and transgenic plants were grown at a constant temperature of 34°C from the day of sowing till the three-leaf stage (approximately four weeks), followed by their growth at 22°C for recovery. We observed 100% germination of both transgenic lines and wild-type (n/n) plants at 34°C. However, seedlings of only transgenic lines survived till the three-leaf stage. The wild type plants died with visible symptoms of heat stress, such as leaf yellowing and drooping (Fig. 2A). During the recovery phase, the transgenic lines fully recovered at 22°C, indicating their improved tolerance to heat stress. Subsequent biochemical and physiological analyses for photosynthetic pigments, PSII activity, cell membrane stability by both electrolyte leakage and lipid peroxidation also supported the enhanced heat-tolerant phenotype of OX lines. In line with the visual observation, we found that under constant Heat Shock (HS) conditions, transgenic plants retained a higher level of chlorophyll pigments (Chl a, Chl b and carotenoids) (Fig. 2B). These plants also showed better photosynthetic yield (YII) (average %), maximum photosynthetic efficiency (Fv/Fm) (average%) and photosynthetic electron transport rate (ETR) related to PSII (average %), than the wild type plants at constant heat stress of 34°C (Fig. 2C). Because the photosynthetic activity is not only dependent on the chloroplastic pigments but also membrane stability, we next checked membrane stability by both electrolyte leakage and lipid peroxidation. We observed that the transgenic lines significantly preserved the membrane stability under heat stress as significantly less lipid peroxidation and lower solute leakage was observed in these plants (average- 24.8%) than their WT counterparts (average- 38.8%) under constant HS condition of 34°C regime.

**Fig. 2.**
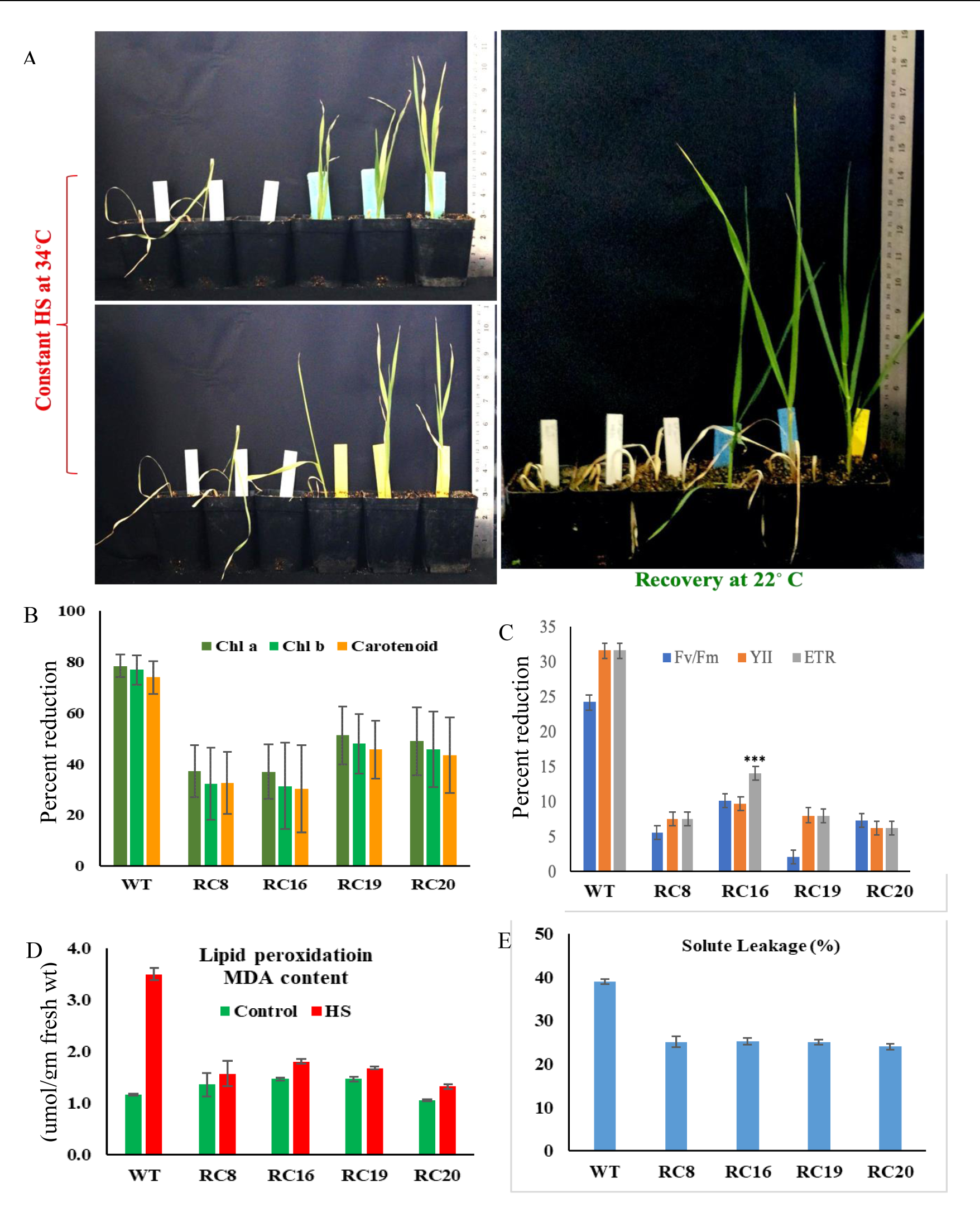
*ZmUbi:HvHsfA6a* plants are more resistant to heat stress A) Phenotypes of wild type and transgenic lines during constant heat stress at 34°C and their recovery at 22°C. B) Estimation of photosynthetic pigments, Chl a, Chl b and carotenoids C) Measurements of PSII activity, including maximum photosynthetic efficiency Fv/Fm, photosynthetic yield YII and electron transport rate. Membrane lipid peroxidation and electrolyte leakage E) of both transgenic and wild type plants. Data represented as the mean values + SD of five biological replicates. *** represents the P value significance < 0.01

### The *ZmUbi:HvHsfA6a* OX plants show lower ROS accumulation and enhanced antioxidative potential

One of the direct harmful effects of heat stress is the enhanced accumulation of various reactive oxygen species and free radicals. Since we observed high heat stress tolerance in the transgenic lines, we next measured the ROS levels along with the activity of detoxifying enzymes such as Super oxide dismutase (SOD), Ascorbate peroxidase (APX), and Catalase (CAT) in these plants. Under optimal control conditions, transgenic and wild-type leaf tissue showed no significant difference in H2O2 and superoxide species accumulation in the NBT and DAB staining. However, wild type plants accumulated more ROS than leaves of RC8 and RC16 transgenic lines under high-temperature conditions (Fig. 3A, 3B). In contrast, the leaves of both transgenic lines plants showed increased activity of all three antioxidant enzymes compared to wild-type plants (Fig. 3C).

**Fig. 3.**
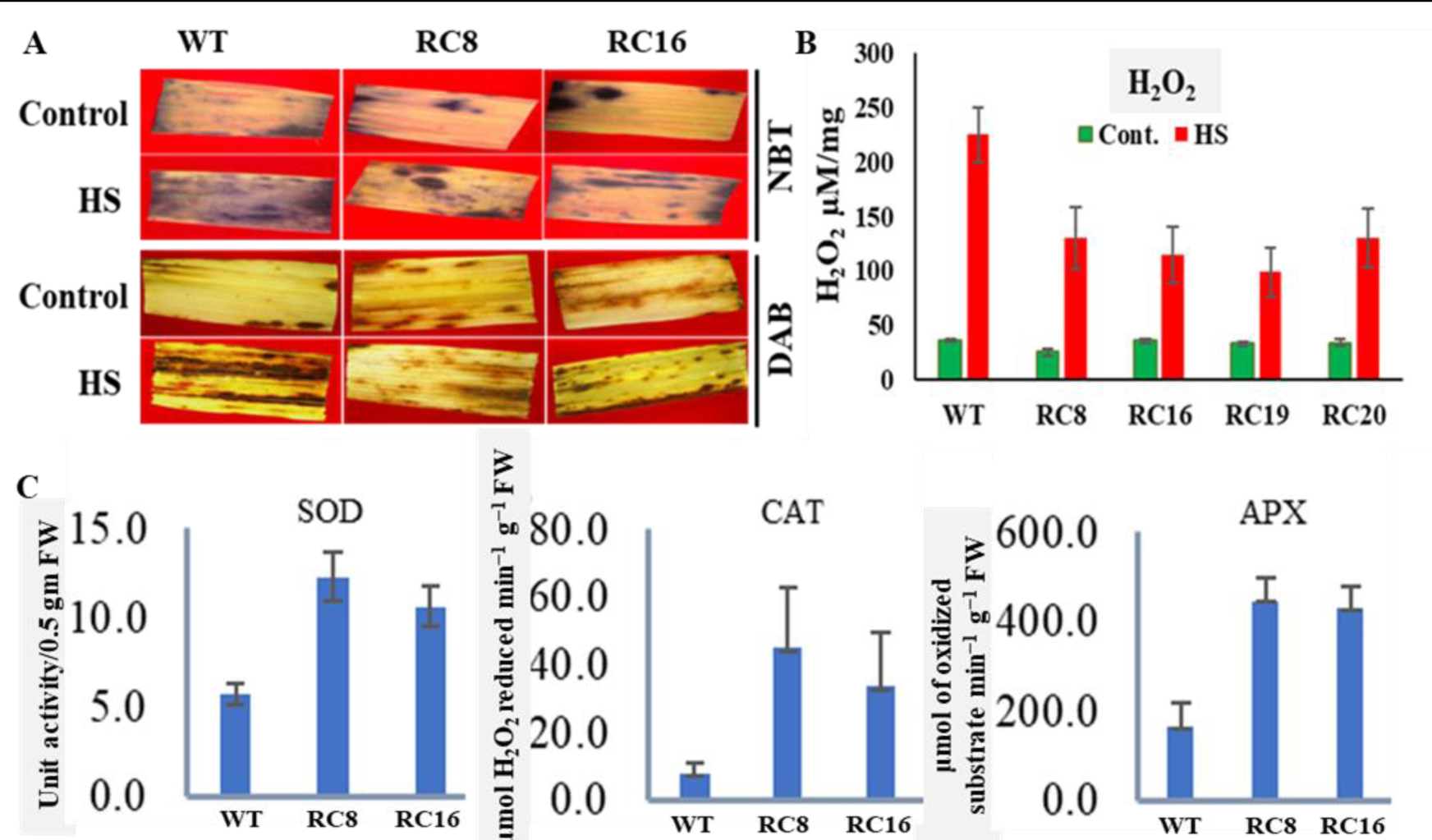
Accumulation of ROS and antioxidative enzymes activity in WT and transgenic lines. A. ROS staining by NBT and DAB stains, B. Measurements of in situ H_2_O_2_ by Amplex red reagent and activity of antioxidant enzymes SOD, CAT and APX in WT and *ZmUbi:HvHsfA6a* plants C, Error bars SD (n=5), P value <0.001

### Transcriptome analysis of *ZmUbi:HvHsfA6a* OX transgenic plants

To understand the molecular basis of the better heat stress tolerance phenotype of the HSFA6a OX plants over their WT counterparts, we performed RNA sequencing of transgenic lines RC8 and RC16 along with two wild-type plants (RCWT1 and RCWT2). Leaf tissues from five-weeks-old plants were used RNA isolation and library preparation. A total of four samples were sequenced on the Illumina platform with a maximum paired-end read length of 151 bp. 80 million high-quality reads were generated for all four samples (Table 1). These reads were mapped on the reference genome of barley downloaded from the Ensemble database (https://plants.ensembl.org/Hordeum_vulgare/Info/Index) using the TopHat pipeline with default parameters. The cufflinks pipeline was used to construct gene models based on TopHat results, and the Cuffdiff pipeline was used for the identification of differentially expressed genes (DEGs). The expression of genes was estimated on the basis of the value of FPKM reads (Fragments Per Kilobase per Million reads) of transcripts.

**Table 1:** Summary of total number of high-quality reads and bases obtained from RNA-Seq of barley

| S. No. | Sample name | Total high-quality reads | Total high-quality bases |
| --- | --- | --- | --- |
| 1. | RC8 and RC16 | 43,092,476 | 3,087,450,574 |
| 2. | RCWT-1 and RCWT-2 | 40,140,844 | 2,978,890,272 |

A principal component analysis of all the obtained transcripts from both wild-type and transgenic plants revealed significant differences in the transcriptomes of transgenic vs WT (Fig. 4A). The differential expression was defined by log_2_ ratio ≥2 of transcripts. In total, 1285 (793 upregulated and 492 downregulated) transcripts were were identified as differentially expressed genes in transgenic lines as compared to wild type lines (Fig. 4B).

**Fig. 4.**
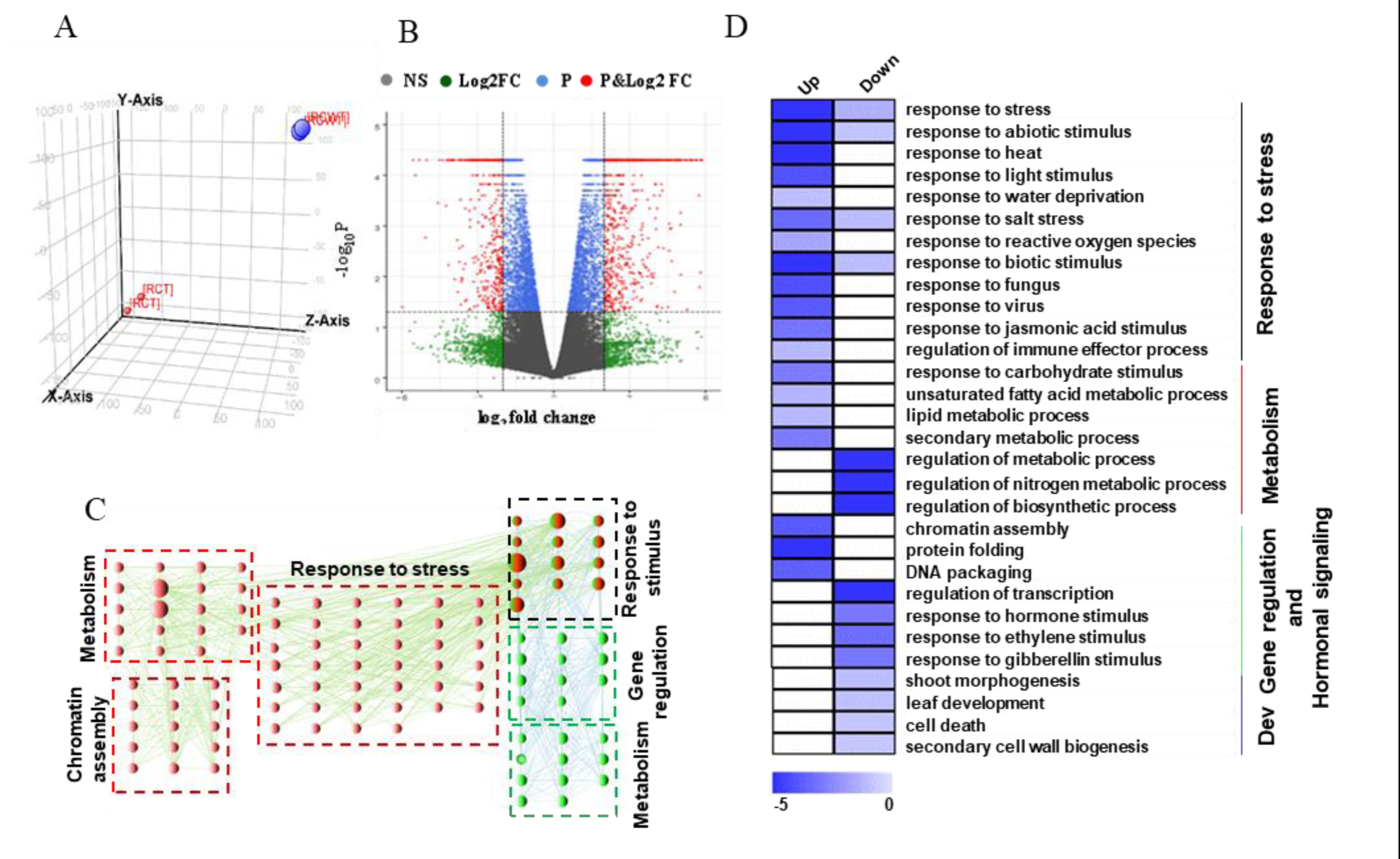
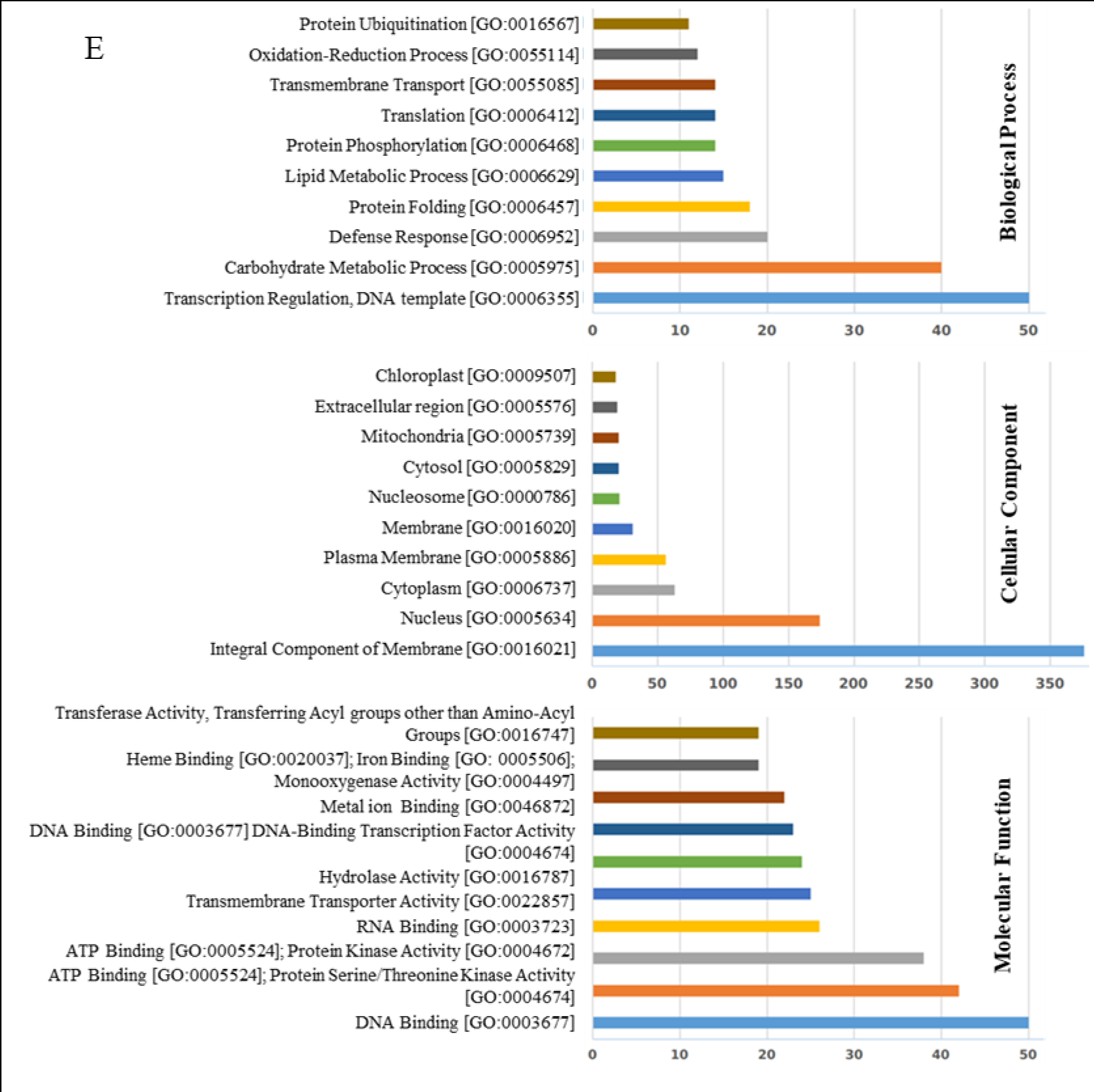
RNA sequencing-based transcriptome analysis of overexpressing transgenic plants compared to wild type. A) PCA of transcriptome of two wild type and two transgenic lines. Red and blue dots indicate transgenic and wild type lines respectively. B) Volcano plots for genes showing differential expression among wild type and transgenic lines. NS-Not Significant, log2FC- transcripts with -2<>2 but not P value significant, P- transcripts with P value < 0.05, log2FC& P: transcripts with log2FC-2<>2 and P value significant.C) Enrichment of GO terms (biological process) between upregulated and downregulated sets of genes are shown via network. Red circles connecting with green thin lines represent sets of enriched GO terms in set of upregulated genes. Likewise, green circles with blue thin lines represent sets of enriched GO terms in set of downregulated genes. Circles with green and red represent enrichment of GO terms in both upregulated and downregulated set of genes. D) Enriched GO terms in set of upregulated and downregulated sets of genes is shown via heatmap. The associated biological processes are labeled at the rightmost side. Scale represent log_10_ corrected *p*-value of enriched GO terms. **E.** Gene ontology analysis of differentially expressed genes among wild type and transgenic lines (*ZmUbi:HvHsfA6a*)

To validate the data generated through RNA-seq, the quantitative RT-PCR analysis of 8 differentially expressed genes of heat shock protein family namely *HvHsp70-2*, *HvHsp70-5*, *HvHsp110-4*, *HvHsp90-2*, *HvHsp90-4*, *HvHsp90-6* and *HvHsp90-7* was performed (Supplementary Fig. S2). We obtained consistent results between qRT-PCR and transcriptome data, which confirms the relaiblity of the data.

Gene Ontology or GO enrichment analysis was utilized to identify the functions of these over- represented transcripts (Fig. 4, C, D, E). These DEGs were categorized based on molecular function, biological process, and cellular components.

This analysis showed that the processes regulating the primary metabolism including photosynthesis, plastid formation, respiration, carbohydrate and lipid metabolism, along with other genes involved in heat stress amelioration, signalling, transcription factors and development are affected in the OX plants. A careful examination of the ten most abundant GO categories in cellular component, biological process and molecular function revealed the important role of membrane organs and processes such as carbohydrate metabolism (photosynthesis and respiration, correlated to chloroplast and mitochondria), nucleic acid metabolism (correlated to nucleus and cytoplasm) and defence related categories (Fig. 4E).

### Genes related to photosynthesis, respiration and chloroplast development are upregulated in transgenic plants

In plants, components of photosynthetic processes such as photosystem II, electron transport chain and enzymes involved in both light and dark reactions are highly vulnerable to heat stress.

In our study, transgenic lines showed upregulation of genes involved in the process of cyclic and non-cyclic electron flow, such as, *FAD/NAD(P)-binding oxidoreductase, ferredoxin-NADP reductase*, regulatory proteins like *thioredoxins*, genes associated with the assembly of PSII and PSI such as reaction centre protein *D1* biogenesis, and *PsbP-domain containing protein 1* (*ppd1*), genes required for proper assembly of Rubisco enzyme and maintenance of its activity including *rubisco large chain*, *small chain*, *rubisco accumulation factor* and *rubisco activase,* respectively. In addition, genes related to the components of electron transport chain, such as, *NADH dehydrogenase complex I*, *Cytochrome c oxidase biogenesis*, and *ubiquinone oxidoreductase*, heme and tetrapyrrole synthesis (e.g. glutamyl*-tRNA reductase-binding protein*, *glutamyl-tRNA(Gln) amidotransferase subunit A*) and chloroplast development related genes (e.g. *plastid division protein* (*PDV*) *PDV1*) showed upregulation in transgenic lines. The details related to abovementioned genes and other upregulated genes such as, gene id, gene name, fold change are given in Supplementary Table S2.

### Upregulation evidenced in stress response and signalling genes

Chaperones are crucial for maintaining cellular protein homeostasis and providing tolerance against heat, drought, oxidative and other abiotic stresses to plants. Here we observed the upregulation of HSP family genes including 5 *HSP70,* 2 *HSP90* and 2 *HSP100*, 13 *DNAJ* chaperonin genes, 5 *ankyrin repeat* family protein, and non-classical chaperones (such as, 3 *protein disulfide isomerase*, 3 *peptide-prolyl cis-trans isomerase* (*PPIases*), and *HSA32)*. Additionally, the upregulation of genes related to hypoxia, salt, and oxidative stress tolerance comprising of *alcohol dehydrogenase*, *aldehyde dehydrogenase*, and *betaine aldehyde dehydrogenase* were found upregulated in *ZmUbi:HvHsfA6a* lines. Besides, several genes involved in scavenging of ROS, such as *Fe-superoxide dismutase*, *catalase*, *glutathione synthetase*, *glutathione peroxidase*, *glutathione-S-transferase* family, and *peroxidases* family gene showed differential expression in *ZmUbi:HvHsfA6a* transgenic lines. The details related to abovementioned genes and other upregulated genes such as, gene id, gene name, fold change are given in Supplementary Table S2

Role of Ca^2+^ ions is well defined in heat stress signalling. *ZmUbi:HvHsfA6a* lines showed differential expression of *calcium dependent protein kinases* (CDPKs), *calmodulins*, *calmodulin-binding proteins*, and *calmodulin-binding transcription activator* (Supplementary Table S2). Similarly, related to kinases, we observed the differential expression of 116 kinase encoding genes (72 upregulated and 44 downregulated), belonging mainly to the family of receptor kinases (RLK), serine/threonine kinases, and leucine rich receptor-like protein kinases (LRR-RLK) (Supplementary Table S2). Finally, *ZmUbi:HvHsfA6a* lines also contained differentially regulated transcription factors which govern the regulation of heat shock response in plants. These TFs include *DREB-element binding protein*, *DREB 1B*, *bZIP* (*HORVU5Hr1G014170*), *ethylene response transcription factor 1* (*ERF1*), 5 heat shock factors (*HvHsfB2c, HvHsfB1, HvHsfA4b*, *HvHsfC1b, HvHsfC2a*), 13 *MYB* family genes, 13 *basic helix-loop-helix* (*bHLH*) family transcription factors which showed both up and downregulation. Further details are given in Supplementary Table S2.

### Transgenic lines showed upregulated fatty acids, amino acids and sugar metabolism-related genes

During heat stress, maintenance of plasma membrane integrity, amino acid metabolism, carbohydrate metabolism, and partitioning of sugar is crucial for survival of plants. In *ZmUbi:HvHsfA6a* lines, we found the differential expression of genes encoding enzyme involved in lipid biosynthesis and signalling processes such as *lipoxygenases I*, *II*, *3-oxoacyl-[acyl-carrier-protein] reductas*e, and *diacylglycerol kinase,* etc. (Supplementary Table S2).

Upregulation of several genes involved in amino acid metabolism which play a key role in establishing cellular homeostasis during heat stress, was observed, capturing higher expression of amino acids like alanine, cysteine, glycine, asparagine, phenylalanine, aspartate, glutamate etc. (Supplementary Table S2). Conversely, related to carbohydrate metabolism, many genes showed differential expression in our overexpressing transgenic lines involved in key processes such as glycolysis (*NADP-dependent glyceraldehyde-3-phosphate dehydrogenase* (*GAPDH*), *ATP-dependent 6-phosphofructokinase (PFK)*, *pyruvate dehydrogenase* (*PDH*) etc.), pentose phosphate pathway (*D-ribulose-5-phosphate-3-epimerase*, *6-phosphogluconolactonase* etc.), citric acid cycle (*malate dehydrogenase*, *aconitase*), and sucrose and starch metabolic processes (*sucrose synthase*, *sucrose-phosphate phosphatase*, *starch synthase*, *acid beta-fructofuranosidase* etc.) (Supplementary Table S2). Upregulation of some sucrose transporter genes, namely *SUT1*, *SUT2*, *SUT7*, *SUT11*, and *bidirectional sucrose transporter N3* involved in partitioning of sugars, were also observed in our overexpressing transgenic lines. Besides, the upregulation of *galactinol synthase* encoding gene which aids in protecting from oxidative damage to plants during heat stress was also unveiled in *ZmUbi:HvHsfA6a* lines. The details related to abovementioned genes and other upregulated genes such as, gene id, gene name, fold change are given in Supplementary Table S2

### Genes related to phytohormone, and secondary metabolism related genes were upregulated in transgenic lines

The phytohormone jasmonic acid is ubiquitous to plant kingdom and plays a key role during biotic and abiotic stresses. We observed the upregulation of the genes involved in its biosynthesis pathway, such as *phospholipase A1-II*, *lipoxygenase, allene oxide synthase*, and *jasmonate O-methyltransferase*. It is well known that secondary metabolism allows plants to combat several adverse environmental conditions, in *ZmUbi:HvHsfA6a* lines we observed the upregulation of genes involved in terpenoid (*terpene synthase, terpene cyclase*) and flavonoid biosynthesis (*dihydroflavanol-4-reductase*) (Supplementary Table S2). The details related to abovementioned genes and other upregulated genes such as, gene id, gene name, fold change are given in Supplementary Table S2.

### Search for targets of HvHsfA6a by ChIP-sequencing analysis

We performed ChIP-sequencing on *ZmUbi:HvHsfA6a* transgenic plants to identify the genome-wide potential targets of HvHsfA6a. For ChIP-Seq, the sonicated DNA before immunoprecipitation was used as negative control input DNA to eliminate non-specific peaks. In this analysis, 6221 HvHsfA6a-specific peaks were identified (Supplementary Table S3). The peak sequences were distributed on all seven chromosomes of the barley genome. Further, the genes associated with these peaks were identified and presented in Supplementary Table S4. Notably, these genes were involved in photosynthetic process, carbohydrate metabolism, chaperones, TFs, fatty acid and amino-acid metabolism and developmental processes (Supplementary Table S4). The analysis of the peak distribution revealed that around one-fifth (20.12%) of the peaks are from putative promoter regions, while another 14 percent belong to other coding regions, such as UTRs and introns (Supplementary Fig S6A). The rest are from distal intergenic regions (65.84%).

GO analysis revealed the enrichment of biological processes including, regulation of transcription, carbohydrate metabolism, primary metabolic processes, defense response, membrane integrity, and transmembrane transport, which suggested the potential role of *HSFA6a* (Supplementary Fig S6B). We then performed a targeted search based on literature for heat stress-specific genes in this data set and identified many motifs and corresponding genes related to heat stress response Table 2). Interestingly, most of the genes whose promoters were enriched in ChIP-Seq analysis were also found upregulated in RNA-Seq analysis. A total of 32 genes were identified which consist of a consensus HSE in their promoter regions. These genes are mainly involved in normal plant growth and HS response such as ethylene responsive, floral development, secondary metabolism, lipid biosynthesis, HSFs etc. could be under the direct regulation of HSFA6a (Supplementary Table S5). However, this could be validated through further experiments.

**Table 2:**
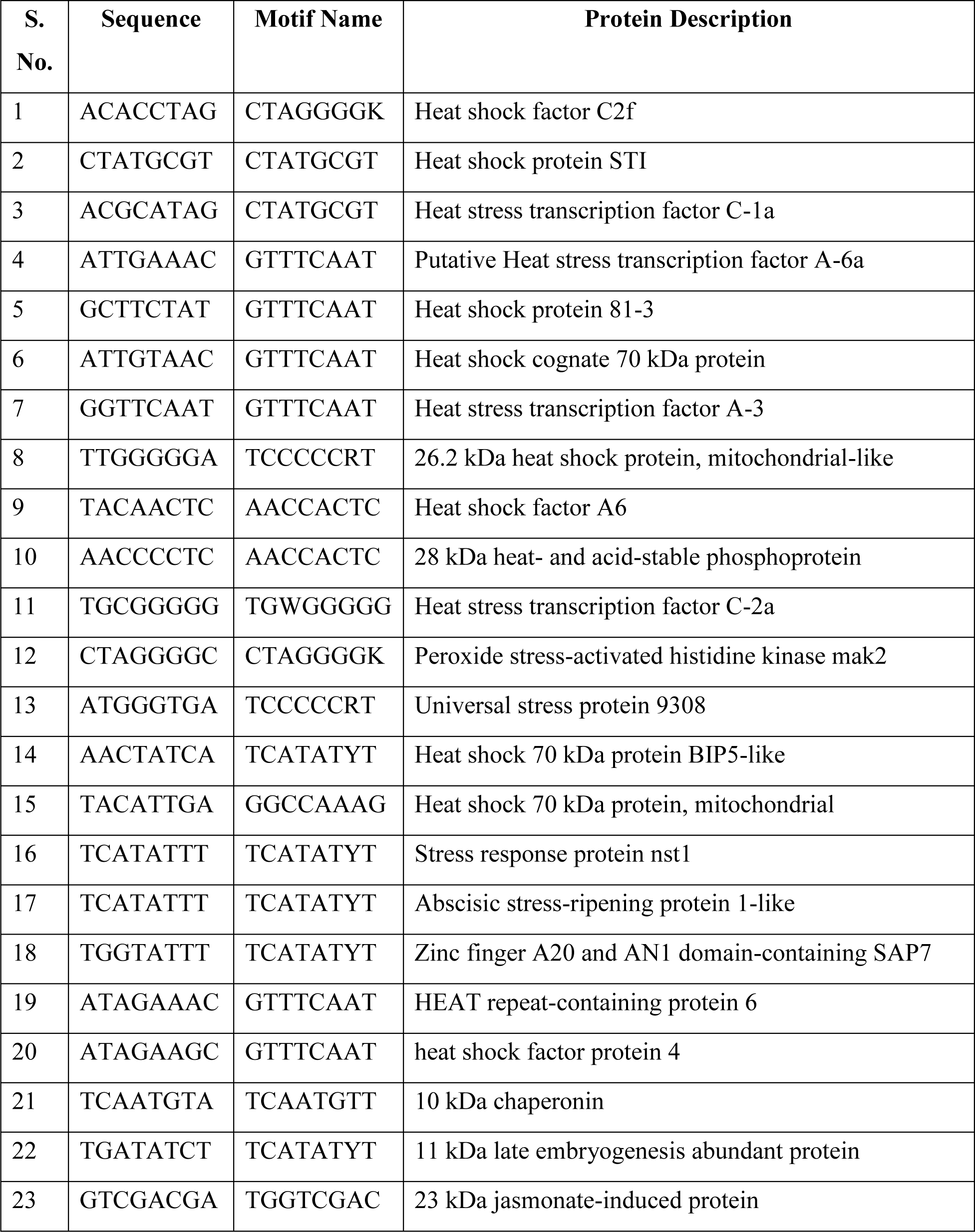
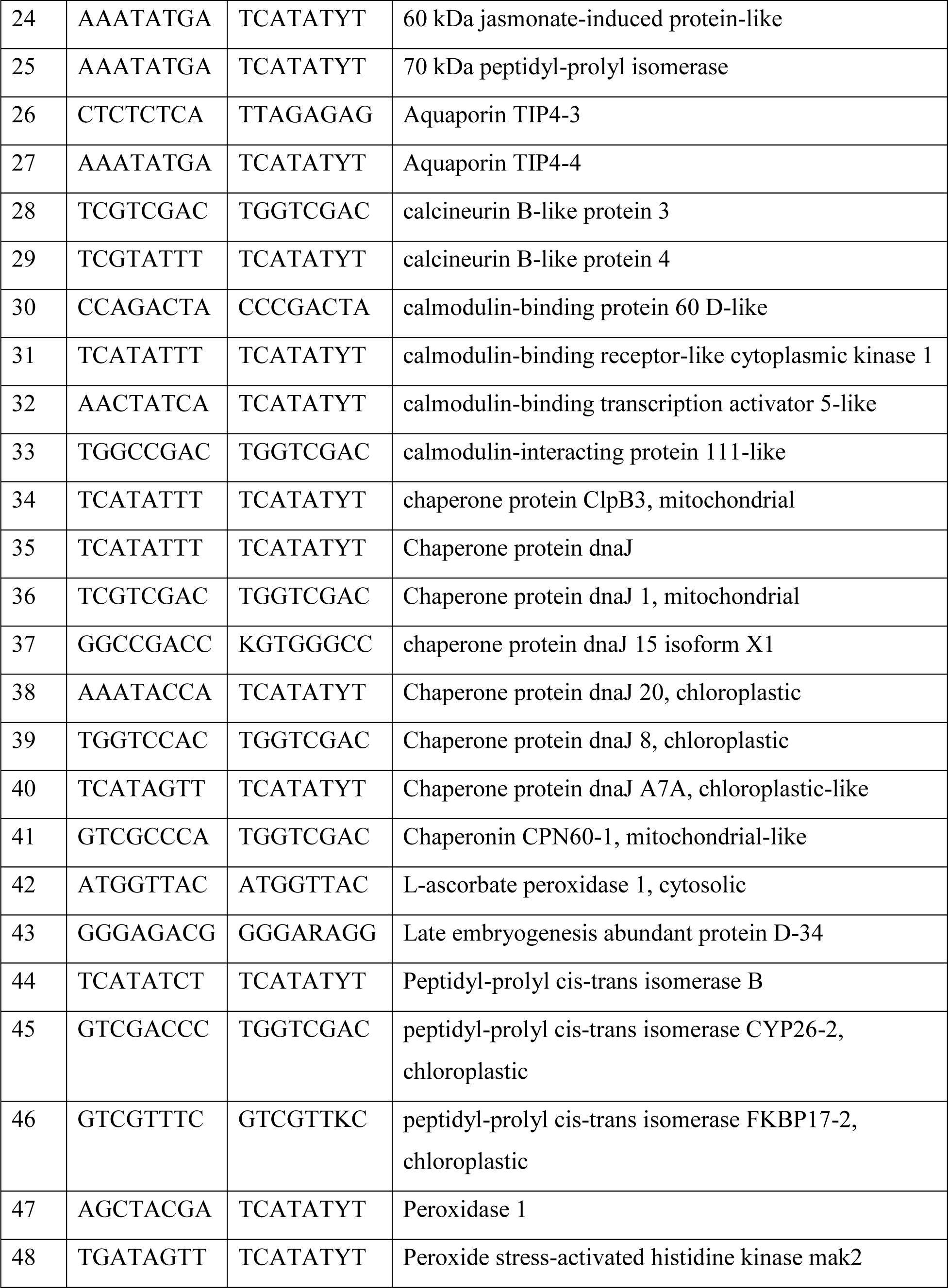
Heat stress related motifs overrepresented in ChIP sequencing data of *ZmUbi:HvHsfA6a* plants

### Transgenic plants showed altered metabolism mimicking heat stress under control and heat stress conditions

Because we found many primary metabolism-related genes were differentially expressed in the OX lines in our transcriptome analysis, we next studied the metabolic profiles of both WT and transgenic plants. GC-MS metabolite profiling of leaf metabolites (wild-type control (WC), wild-type heat stress treated (WH), transgenic control (TC), and transgenic heat stress-treated (TH), identified a total of 39 differentially accumulating primary metabolites across all the samples (Table 3). These metabolites mainly include amino acids, organic acids, vitamin, secondary metabolites, sugar and sugar alcohols. One-way ANOVA *p*-value < 0.05 was used to compare the metabolites from WC, WH, TC and TH and Log_2_ fold change values between different sample types are given in Table 4. To check the contribution of these metabolites in distinguishing the samples, a supervised partial least squares discriminant analysis (PLS-DA), was performed. The 2D score plots obtained from the PLS-DA test showed that the metabolites of WC, WH, TC and TH samples did not overlap with each other, indicating an altered state of metabolite levels (Fig. 5A). This separation of four treatment types indicates a different metabolite level in each treatment type. The first two components (component 1 and component 2) of PLS-DA accounted for 67.1% and 26.9% variance among samples, respectively. The influence of the metabolites (independent variable) in explaining the dependent variable (treatment type) was estimated by VIP scores where metabolites with VIP score (≥1) were considered as important metabolites to play crucial roles in distinguishing the treatment types (Fig. 5B). The heatmap for the total metabolites from all the sample types is shown in Fig. 5C. Fifteen differential marker metabolites namely, palmitic acid, talose, D-fructose, myo-inositol, oxalic acid, sucrose, D-trehalose, D-glucose, glycine, D-xylose, L-threonine, L-aspartic acid, D-threitol, malic acid, and D-galactose were detected based on VIP scores (Supplementary Table S6, 7). It is worth to not that in wild-type plants, the amount of important sugars is decreased upon heat stress ( Table 4 and Fig. 5C ). On the other hand, no such decrease in sugar levels was observed in the HvHsfA6a overexpression plants; which still maintained a positive fold change correlating to a sustained supply possibly from higher photosynthetic activity shown by these plants even under heat stress condition. In addition, metabolites such as p-coumaric acid, caffeic acid, shikimic acid, and chlorogenic acid, and known to be involved in both abiotic and biotic stress also showed enhanced accumulation in transgenic lines under heat stress as compared to their lower accumulation in wild-type plants.

**Fig. 5.**
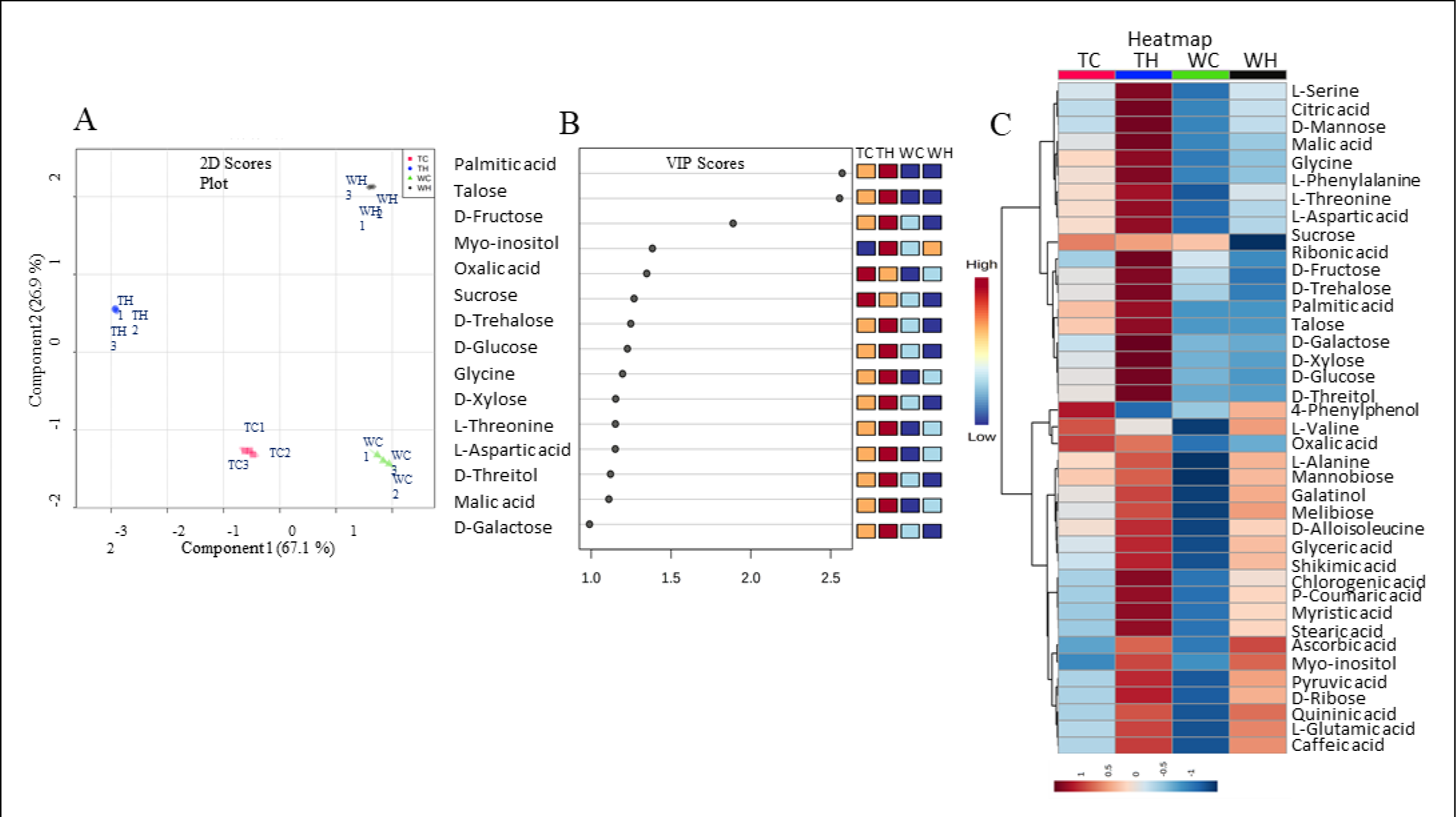
Transgenic plants showed higher accumulation of heat stress related metabolites under both control and heat stress conditions. A) 2D scores plot (PLS-DA), B) Important features (PLS-DA, VIP scores) and C) Heatmap analyses of GC- MS data by using Metaboanalyst 3.0. In scores plot, metabolites from wild type control (WC), wild type heat stress (WH), transgenic control (TC) and transgenic heat stress (TH) leaves didn’t overlap with each other, indicating an altered state of metabolite levels.

**Table 3.**
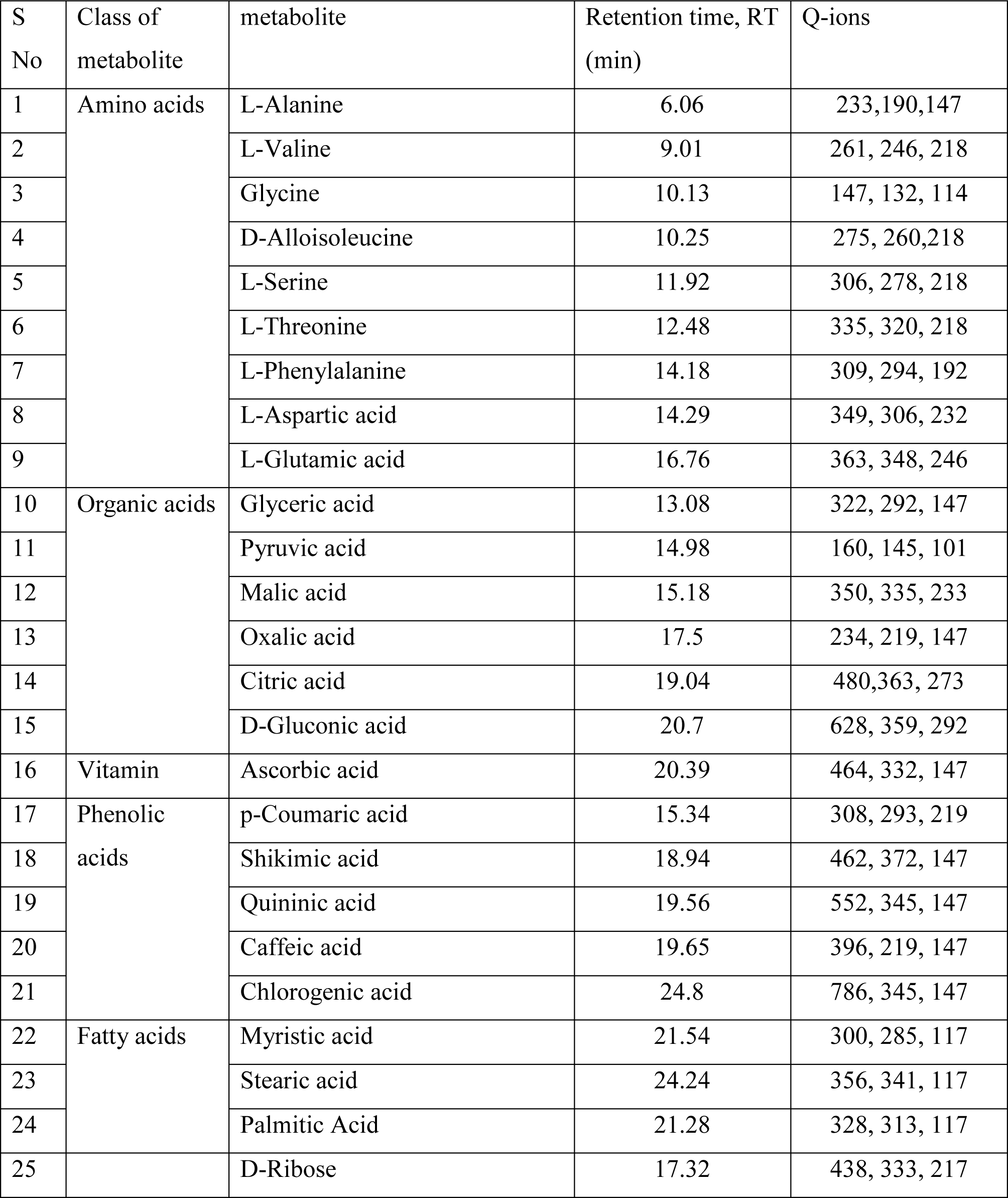
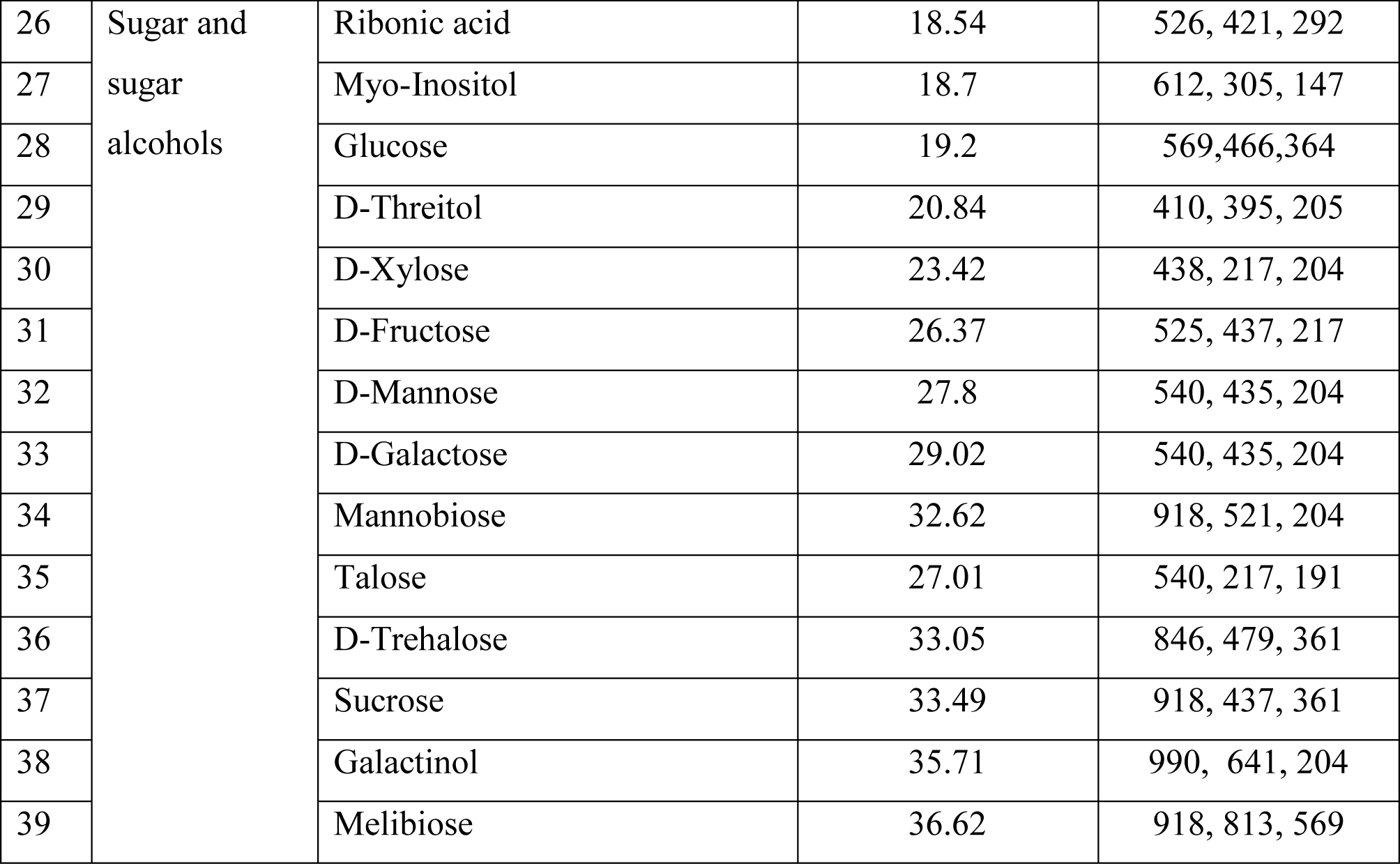
GC-MS-based metabolite profile of wild type and transgenic wheat leaf under heat stress conditions. Separation was carried out on HP5-MS GC column. Metabolites were identified based on matching with RT and mass spectrum of standard reference compounds.

| S No | Class of metabolite | metabolite | Retention time, RT (min) | Q-ions |
| --- | --- | --- | --- | --- |
| 1 | Amino acids | L-Alanine | 6.06 | 233,190,147 |
| 2 |  | L-Valine | 9.01 | 261, 246, 218 |
| 3 |  | Glycine | 10.13 | 147, 132, 114 |
| 4 |  | D-Alloisoleucine | 10.25 | 275, 260,218 |
| 5 |  | L-Serine | 11.92 | 306, 278, 218 |
| 6 |  | L-Threonine | 12.48 | 335, 320, 218 |
| 7 |  | L-Phenylalanine | 14.18 | 309, 294, 192 |
| 8 |  | L-Aspartic acid | 14.29 | 349, 306, 232 |
| 9 |  | L-Glutamic acid | 16.76 | 363, 348, 246 |
| 10 | Organic acids | Glyceric acid | 13.08 | 322, 292, 147 |
| 11 |  | Pyruvic acid | 14.98 | 160, 145, 101 |
| 12 |  | Malic acid | 15.18 | 350, 335, 233 |
| 13 |  | Oxalic acid | 17.5 | 234, 219, 147 |
| 14 |  | Citric acid | 19.04 | 480,363, 273 |
| 15 |  | D-Gluconic acid | 20.7 | 628, 359, 292 |
| 16 | Vitamin | Ascorbic acid | 20.39 | 464, 332, 147 |
| 17 | Phenolic acids | p-Coumaric acid | 15.34 | 308, 293, 219 |
| 18 |  | Shikimic acid | 18.94 | 462, 372, 147 |
| 19 |  | Quinic acid | 19.56 | 552, 345, 147 |
| 20 |  | Caffeic acid | 19.65 | 396, 219, 147 |
| 21 |  | Chlorogenic acid | 24.8 | 786, 345, 147 |
| 22 | Fatty acids | Myristic acid | 21.54 | 300, 285, 117 |
| 23 |  | Stearic acid | 24.24 | 356, 341, 117 |
| 24 |  | Palmitic Acid | 21.28 | 328, 313, 117 |
| 25 |  | D-Ribose | 17.32 | 438, 333, 217 |
| 26 | Sugar and<br>sugar<br>alcohols | Ribonic acid | 18.54 | 526, 421, 292 |
| 27 |  | Myo-Inositol | 18.7 | 612, 305, 147 |
| 28 |  | Glucose | 19.2 | 569,466,364 |
| 29 |  | D-Threitol | 20.84 | 410, 395, 205 |
| 30 |  | D-Xylose | 23.42 | 438, 217, 204 |
| 31 |  | D-Fructose | 26.37 | 525, 437, 217 |
| 32 |  | D-Mannose | 27.8 | 540, 435, 204 |
| 33 |  | D-Galactose | 29.02 | 540, 435, 204 |
| 34 |  | Mannobiose | 32.62 | 918, 521, 204 |
| 35 |  | Talose | 27.01 | 540, 217, 191 |
| 36 |  | D-Trehalose | 33.05 | 846, 479, 361 |
| 37 |  | Sucrose | 33.49 | 918, 437, 361 |
| 38 |  | Galactinol | 35.71 | 990, 641, 204 |
| 39 |  | Melibiose | 36.62 | 918, 813, 569 |

**Table 4:** Fold changes of metabolites in the wild type control (WC), wild type heat stressed (WH), transgenic control (TC) and transgenic heat-stressed (TH) lines of wheat. Fold change was calculated using relative peak area data of each metabolite from three biological replicates obtained through GC-MS.

| Metabolite | Log<sub>2</sub> Fold Change (FC) |  |  |  |
| --- | --- | --- | --- | --- |
|  | FC(WH/WC) | FC(TH/TC) | FC(TH/WC) | FC(TH/WH) |
| L-Alanine | 1.33 | 0.59 | 1.66 | 0.33 |
| L-Valine | 1.35 | 1.10 | 3.69 | 2.34 |
| Glycine | 0.34 | 0.39 | 1.49 | 1.15 |
| D-Alloisoleucine | 0.25 | 1.71 | 1.94 | 1.69 |
| L-Serine | 0.89 | 0.99 | 2.91 | 2.01 |
| L-Threonine | 0.72 | 1.09 | 2.21 | 1.49 |
| L-Phenylalanine | 0.92 | 0.91 | 2.04 | 1.12 |
| L-Aspartic acid | 0.17 | -0.47 | 0.19 | NS (0.01) |
| L-Glutamic acid | 0.93 | -0.45 | NS (0.02) | -0.91 |
| Glyceric acid | 1.47 | 1.09 | 2.15 | 0.67 |
| Pyruvic acid | 1.32 | 1.10 | 1.77 | 0.45 |
| Malic acid | 0.46 | 0.79 | 2.30 | 1.84 |
| Oxalic acid | 0.15 | 0.41 | 0.69 | 0.54 |
| Citric acid | 0.45 | 1.60 | 1.52 | 1.07 |
| D-Gluconic acid | 0.18 | 0.56 | 1.00 | 0.81 |
| Ascorbic acid | 1.46 | 0.97 | 2.14 | 0.68 |
| p-Coumaric acid | 0.83 | 1.08 | 1.56 | 0.73 |
| Shikimic acid | 1.19 | 0.84 | 1.84 | 0.65 |
| Quinic acid | 0.97 | 1.11 | 1.03 | 0.06 |
| Caffeic acid | 0.92 | 0.32 | 1.14 | 0.22 |
| Chlorogenic acid | 0.77 | 0.63 | 1.66 | 0.89 |
| Myristic acid | 0.81 | 1.15 | 1.56 | 0.75 |
| Stearic acid | 0.82 | 1.21 | 1.62 | 0.80 |
| Palmitic Acid | 1.08 | 1.99 | 3.12 | 2.88 |
| D-Ribose | 0.57 | 1.24 | 1.15 | 0.58 |
| Ribonic acid | -0.30 | 0.46 | 0.66 | 0.96 |
| Myo-Inositol | 2.79 | 3.12 | 3.04 | 0.25 |
| Glucose | -0.14 | 1.12 | 1.58 | 1.71 |
| D-Threitol | **NS (-0.04)** | 0.93 | 1.36 | 1.41 |
| D-Xylose | -0.09 | 0.69 | 1.64 | 1.73 |
| D-Fructose | -1.17 | 2.24 | 2.65 | 3.81 |
| D-Mannose | 0.40 | 1.01 | 1.42 | 1.01 |
| D-Galactose | **NS (-0.06)** | 0.75 | 1.65 | 1.72 |
| Mannobiose* | ND | 0.45 | ND | 0.62 |
| Talose** | ND | 2.30 | ND | ND |
| D-Trehalose | -0.40 | 0.58 | 1.24 | 1.65 |
| Sucrose | -0.62 | 0.13 | NS (-0.07) | 0.69 |
| Galactinol* | ND | 1.60 | ND | 0.85 |
| Melibiose* | ND | 1.74 | ND | 0.68 |

Fold changes were calculated using the formula $\text{Log}_2(\text{WH/WC})$ , $\text{Log}_2(\text{TH/TC})$ , $\text{Log}_2(\text{TH/WC})$ and $\text{log}_2(\text{TH/WH})$ . ND: not detected. \* not detected in WC; \*\* not detected in WC and WH. $P < 0.05$ ; NS: Non-significant.

### Pathway analysis reveals markers of heat stress tolerance of a limited set of metabolic pathways

Further, pathway analyses (Fig. 6, A and B) by Metaboanalyst 3.0 using KEGG pathway and rice plant databases, revealed differential metabolite markers of heat stress tolerance mainly altered the following metabolic pathways: (1) alteration in the galactose metabolism (2) alteration in starch and sucrose metabolism which induces sucrose synthesis (3) alteration in fatty acid metabolism which induces fatty acid synthesis (4) alteration in tricarboxylic acid cycle (TCA) metabolism which induces organic acid biosynthesis (5) alteration in carbon metabolism of photosynthesis (6) alteration in fructose metabolism (7) alteration in amino acid metabolism (Fig. 6, Supplementary Fig. S7). It is also worth mentioning here that gene related to these pathway and metabolites are also upregulated in transgenic plants (Supplementary Fig. S3-S5).

**Fig. 6.**
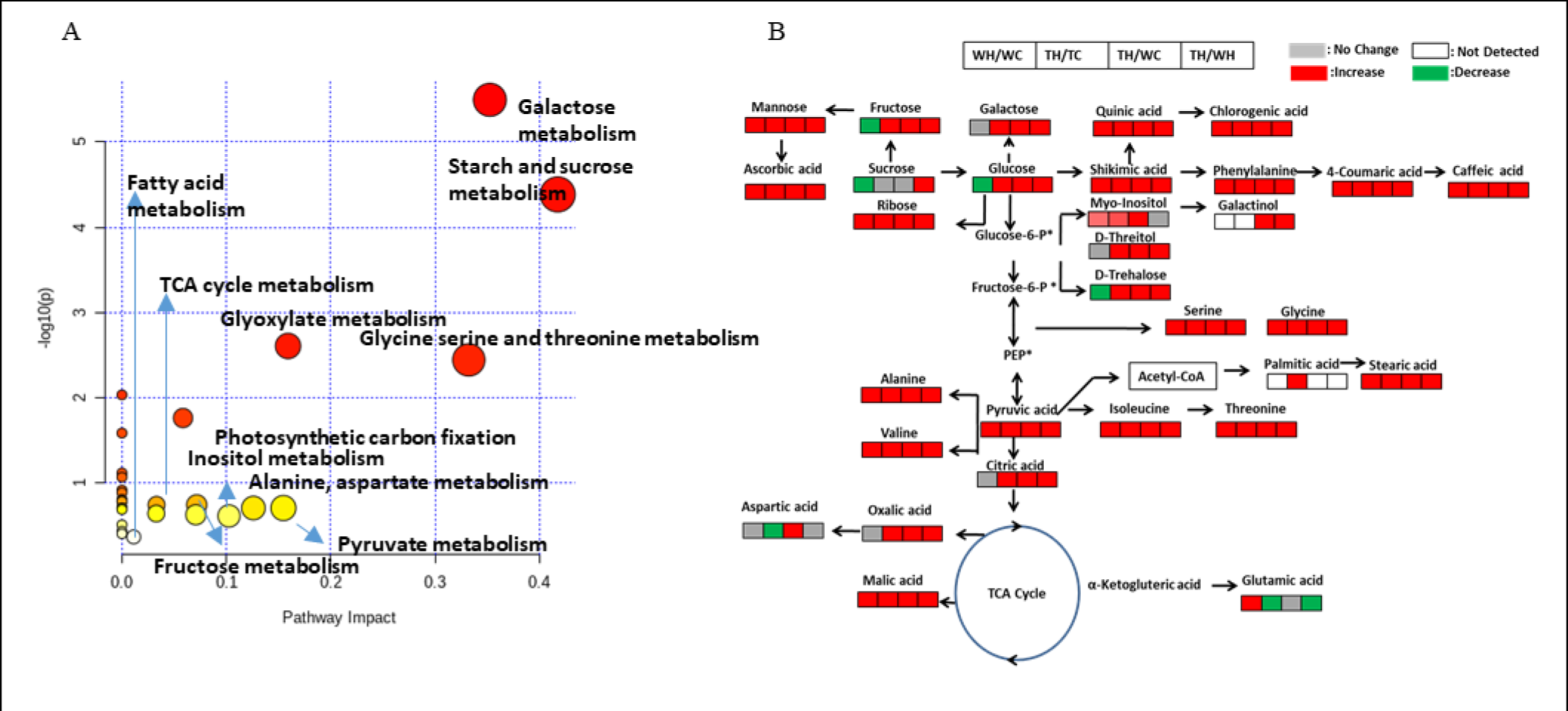
Pathway analysis reveals markers of heat stress tolerance of a limited set of metabolic pathways. A) Pathway analysis based on metabolite markers detected control and heat-treated wild type and transgenic barley leaves. B) Metabolic alterations changes among wild type control (WC), wild type heat stress (WH), transgenic control (TC) and transgenic heat stress (TH) wheat leaf samples from fold change analyses. The proposed metabolic pathways were based on web-based metabolic pathway databases and the literature search.

## Discussion

Heat stress and other environmental challenges disturb the overall homeostasis of plant cells. In response to these adversities, transcriptional reprogramming of several stress responsive genes and metabolic alterations occurs in the cell. The expression of these genes and their underlying regulatory networks which eventually confer tolerance towards various abiotic stresses to plants are known to be under the tight regulation of class A HSFs (Ogawa et al., 2007, Chan-Schaminet et al., 2009, Banti et al., 2010, Scharf et al., 2012, Wang et al., 2014, Xue et al., 2014). Here, we investigated the role of HvHsfA6a of class A subfamily in conferring thermotolerance to barley plants through over expression, and furthermore identified the underlying stress responsive genes and metabolites which directly or indirectly provide thermotolerance. By utilizing *in silico* analysis based on motif composition, and later subcellular localization assay, we notably discovered HvHsfA6a as a nucleus residing transcription factor under both controlled and heat shock conditions. Moreover, qRT-PCR based analysis revealed its early (within 10 minutes of heat stress exposure) and high relative expression as compared to other HSFs present in barley (Mishra et al., 2020). This further makes this protein as key putative candidate regulating HSR in barley.

Heat stress primarily affects the photosynthetic and respiratory reactions, and thereby the carbon assimilation processes (Song et al., 2014, Bita and Gerats, 2013). It adversely affects the photosynthetic processes by disturbing the structural organization of photosystems and flow of electrons, causing denaturation of vital enzymes and eventually leads ROS production (Salvucci and Crafts-Brander et al., 2004).

In our study, during heat stress, *HvHSFA6a* overexpressing lines showed improved photosynthesis as compared to control plants. These findings were supported by transcriptome analysis of these lines, where a number of genes showed higher upregulation and were related to structural and functional organisation of photosynthetic apparatus (Supplementary Table 2). These genes include *high chlorophyll fluorescence phenotype 173* (*hcf173*) gene related to biogenesis of D1 protein, which is crucial for maintenance of higher abundance of other PSII proteins under heat stress (Chen et al. 2020, Schult et al. 2007). Related to the assembly of PSI, we found the upregulation of *ppd1* gene. Given the mutants of its orthologues gene in Arabidopsis showed abolished PSI activity (Liu et al., 2012), Furthermore, we found upregulation of *FNR* (*ferredoxin-NADP reductase*) and *thioredoxins* in our transgenic lines, which are crucial for protecting chlorophyll degradation during heat stress (Lin et al., 2013). During carbon assimilation process, rubisco activase (RCA) regulates the activity of Rubisco (Ribulose-1,5-bisphosphate carboxylase/oxygenase) by promoting carbomylation of CO_2_ (Spreitzer and Salvucci 2002) upregulated expression of RCA is correlated with the higher plant productivity during heat stress in crop plants such as wheat, Arabidopsis, and maize (Ristic et al., 2009). Upregulation of *RCA* and *RAF1* genes in our transgenic lines along with other abovementioned genes further supports the better photosynthetic activity of these lines. Taken together, these findings suggest that *HvHsfA6a* may be vital candidate regulating the photosynthetic activity of plant in an assertive manner.

Photosynthetic reactions fix carbon in form of sucrose as a final product. Heat stress drastically affects the sucrose metabolism by inhibiting the activity of enzymes involved in sucrose synthesis and catabolism, thus disturbing overall energy metabolism of plants (Ruan et al., 2010, Vasseur et al., 2011, Ruan, 2014). Whereas high availability of carbohydrates such as, sucrose and glucose during heat stress has been associated with the heat tolerant trait in creeping bentgrass (Liu and Huang, 2000). In our study, the upregulation of genes encoding enzymes involved in the synthesis of sucrose such as, *sucrose phosphate synthase* and *sucrose synthase* which convert sucrose into its metabolizable form, i.e. glucose and fructose were found to be upregulated. Metabolomic analysis of these transgenic lines further confirmed the higher accumulation of sugars such as sucrose and galactinol under both normal and heat stressed conditions. Apart from being involved in energy metabolism, increased accumulation of sucrose were found associated with thermopriming, as it acts as primary signalling molecule during early heat stress in Arabidopsis (Serrano et al. 2019). Whereas, galactinol protects from oxidative damage along with maintaining thermomemory in Arabidopsis plants upon heat priming (Serrano et al. 2019, Song et al. 2016). Similarly, related to carbohydrate metabolism in transgenic lines we found the upregulation of genes involved in glycolysis and TCA cycle (Supplementary Table 2) and higher accumulation of their intermediates upon metabolomic study such as pyruvic acid, oxalic acid and malic acid. We further observed the accumulation of amino acids such as alanine, valine, serine which are known to be derived from TCA cycle intermediates such as pyruvate and oxaloacetate and upregulation of genes related to their biosynthesis. Previously Kaplan et al. 2004 showed the accumulation of these amino acids a HSR response in *Arabidopsis* plants. In our study the accumulation of shikimic acid was also found which act as precursor of aromatic amino acids such as phenylalanine and tyrosine. Serrano et al. (2019) found the accumulation of shikimate in heat primed plants as compared to non-primed Arabidopsis plants. Taken together, these results suggested that HvHsfA6a takes part in transcripto-metabolic reprogramming leading to thermopriming in transgenic barley plants (Fig. 5,6 and Table 3, 4).

The expression of heat shock proteins is regulated by HSFs during heat stress (Yoshida et al., 2011, Scharf et al., 2012, Wang et al. 2017). In the overexpression lines, upregulating transcripts of heat shock protein family genes (sHSP, *HSP70*, *90* and *100*), chaperonins and nonclassical chaperones like *protein disulfide isomerase* (*PDI*) and their motif enrichment upon ChIP seq signified that the expression of these HSPs might be potentially under the direct regulation of *HvHsfA6a*. These HSPs not only aid in maintaining the structure, conformation and folding of heat labile protein but also involve in providing acquired thermotolerance (Xue et al. 2014). It has already shown that the accumulation of HSPs in heat primed Arabidopsis, wheat and rice plants, contributed towards enhanced thermotolerance (Wang et al., 2016, Jagadish et al. 2010, Thirumalaikumar et al. 2021).

Furthermore, we observed the upregulation of *protein disulfide isomerase* (*PDI*) genes, which potentially function in maintaining the activity of ER-specific proteins during stressed conditions (Kamauchi et al., 2005, Ali and Mutus, 2014). Interestingly, we found the upregulation of a heat shock protein *HSA32* which has been reported to be induced by HSFA2 for maintaining heat stress memory in rice and Arabidopsis plants (Charng et al., 2006, Lin et al., 2014, Baurle, 2016).Also, In our RNA-Seq dataset of *ZmUbi:HvHsfA6a* lines we observed the upregulation of two FKBP-type *peptide-prolyl cis-trans isomerases* genes, reported as orthologues of Arabidopsis *ROF1* (*FKBP62*). These gene in Arabidopsis is required for establishing HS memory by interacting with Hsp90.1, to maintain HSFA2 in active state (Meiri and Breiman, 2009). These results suggested that *HvHsfA6a* may be preparing plants for any upcoming heat stress exposures, through accumulating chaperones and other memory related genes for maintaining protein homeostasis in barley.

The excessive ROS production due to heat and other abiotic stresses causes peroxidation of cell membrane lipids, impede photosynthesis, respiration, and other cellular processes (Mittler et al. 2011, Choudhary et al. 2017). ROS scavenging is majorly aided by both enzymatic and a non-enzymatic antioxidant pathways (Choudhary et al., 2017, Mittler et al., 2004). Here, our transgenic lines showed increased activity of commonly known enzymatic ROS scavengers such as SOD, APX and CAT (Fig. 3B), accompanied by upregulation of genes associated with antioxidant mechanisms (Supplementary Table 2), and accumulation of nonenzymatic antioxidants such as ascorbic acid as compared to wild type plants. Additionally, heat stress affects the membrane integrity by inducing membrane lipolysis, essential to govern vital cellular functions. In addition of showing upregulation of fatty acid biosynthesis-related genes and accumulation of membrane lipids such as stearic acid and palmitic acid. Our transgenic lines showed higher membrane stability during heat stress to that of wild type control plants, which was confirmed through measuring MDA content and electrolyte leakage. Previously, synthesis of saturated fatty acids such as palmitic acid and an increase in saturated to unsaturated fatty acid ratio has been associated with the heat tolerance in Arabidopsis, tall fescue, soybean and many other plants respectively (Narayanan et al., 2020, Hu et al., 2018, Higashi et al., 2015). Hence, our results depicted the mode of HSR regulation by *HvHsfA6a* in barley during oxidative stress, imposed by heat stress.

Moreover, heat stress induced the alterations in fluidity of plasma membrane and transmits signal of heat stress through opening of Ca +2 channels (Saidi et al., 2009, Lenzoni and Knight, 2019).This increase in flux of Ca +2 ions evokes a heat shock response through interacting with downstream targets such as calmodulins, Calcinurin B-like (CBL) proteins, calcium dependent protein kinases, Calcium-binding EF-hand family proteins, and mediate signalling in response to Ca + (Shi et al., 2018, Lenzoni and Knight, 2019). Wang and Huang, (2017) reported that in tall fescue, HsfA2c provides thermotolerance through calcium-mediated signalling. Here, we also observed the upregulation of calcium binding EF-hand family, calmodulin, CDPKs, and calcinurins in overexpressing *ZmUbi:HvHsfA6a* lines. This implicates that HvHsfA6a of barley could also confer thermotolerance through calcium-mediated signalling.

Given HSFs are the terminal components of heat stress signalling, interestingly, we observed the upregulation of one class A, and two class B HSFs, namely, *HvHsfA4b*, *HvHsfB2c*, and *HvHsfB1* respectively. Among these, *HvHsfA4b* has been found to have regulatory roles during late seed development (Reddy et al., 2014), whereas *HvHsfB2c* acts as the regulator of several sHsps having role in seed development, and confer desiccation tolerance to barley (Reddy et al., 2014). Conversely, *HvHsfB1* showed high upregulation during heat stress in barley (Reddy et al., 2014). These TFs may be directly regulated by HSFA6a, as we found a consensus heat shock element in their promoter region. Together with this, differential regulation of other transcription factors family genes such as, *WRKY*, *MYB*, *DREBP-1B*, *ERF1*, *bHLH,* and *bZIP* evidenced in *HvHsfA6a* overexpressing lines of current study, known to regulate various vital cellular pathways including biotic and abiotic stress specific responses in plants (Katiyar et al., 2012, Yang et al., 2012 and Mizoi et al., 2012). These could be under direct or indirect regulation of HvHSFA6a, although we could not find any HSE upon motif analysis through ChIp-Seq. Previously, it was confirmed that HSFA7b of Arabidopsis regulated salt tolerance by binding to E-box like motif (Zhang et al. 2019). Thus, we speculate that key *HvHsfA6a* may co-ordinately regulate the complex crosstalk between the gene networks to provide HS response in barley.

Phytohormone jasmonic acid has been widely associated with various biotic and abiotic defence responses in plants and recent studies describe role of JA in rice and Arabidopsis during heat and combination of heat and high light stress, respectively (Cohen and Leach, 2019, and Balfagon et al., 2019). It is noteworthy, that in our study, *phospholipase A1-II*, *lipoxygenase, allene oxide synthase*, and *jasmonate O-methyltransferase* genes involved in biosynthetic and regulatory pathways of jasmonic acid were found upregulated. This together suggests that *HvHSFA6a* may be the regulator of jasmonic acid based responses during heat stress in barley, however, this is the subject of future studies.

Together, our data suggest that transgenic plants overexpressing *ZmUbi:HvHsfA6a* show priming effects for heat stress as they accumulated metabolites already in control conditions which are otherwise accumulated in wild type plants under heat stress conditions (Fig 5 and Table 3, 4). This is also correlated well with the global transcriptomics and ChIP sequencing analysis, where genes related to biosynthesis of these metabolites are found already upregulated in transgenic *ZmUbi:HvHsfA6a* plants under control conditions.

## Conclusions

Alleviating yield losses in cereal plants caused by climate change (including rising temperature) create an urgent need for characterization of candidate genes exhibiting the potential to provide tolerance to crop plants against heat stress. HSFs are the key target genes which function as transcriptional regulators of several abiotic stress responsive genes. Therefore, we developed transgenic lines of barley overexpressing *HvHsfA6a* under *ZmUbiI* promoter followed by their transcriptome, ChIP sequencing and metabolome analysis.

Transcriptome and ChIP-seq analysis of the resulting transgenic lines unveiled possible potential underlying mechanisms, especially under the transcriptional regulation of HvHsfA6a, and its co- ordinating putative targets involved in providing thermotolerance to barley plants. Moreover, dissecting transgenic and wild type plants’ metabolome under normal and heat stressed conditions through GC-MS revealed the alterations in vital metabolic fluxes. Eventually, these results revealed the differential accumulation of metabolites which aids plants to combat heat stress. Here, we further speculate that overexpression of *HvHSFA6a* prepares transgenic plants for future heat stress by transcript-metabolomic reprogramming of genes and metabolites which are mainly related to heat stress priming and memory.

Thus, our results effectively captured insights for functional genomics and metabolomics of heat stress under the influence of *HvHsfA6a* transcription factor in barley. Based on outputs, we proposed that *HvHsfA6a* could be one of the potential target candidate gene for generating high temperature resilient plants of Triticeae tribe, by utilizing the biotechnological and molecular breeding approaches. Although, further experiments could be done for further comprehensive study such as yeast one hybrid assays and comparison of transcriptome and metabolome of heat primed and non-primed barley plants.

Fold changes were calculated using the formula Log_2_ (WH/WC), Log_2_(TH/TC), Log_2_(TH/WC) and log_2_(TH/WH). ND: not detected. * not detected in WC; ** not detected in WC and WH. P < 0.05; NS: Non-significant.

## Material and Methods

### Plant material growth conditions and heat treatment

For cloning of *HvHSFA6a* gene, seeds of cultivar Golden Promise were sown in germination trays containing mixture of soil and soilrite (1:1) under controlled conditions in greenhouse 18/16 ^◦^C (day/night) temperature and 16- hour photoperiod with light intensity 136 μmol/m2/s photon flux density. Two-weeks-old plantlets were subjected to heat stress at 42 ^◦^C for different durations i.e. for 15, 30, 45, 60 and 120 minutes in a growth chamber. The leaf samples were harvested at every time point and snap freezed in liquid nitrogen. Similarly, for expression analysis of *HvHsfA6a* gene, heat stress was given to the two-weeks-old seedling for 2 hour at 42 ^◦^C under the same growth conditions.

For tissue culture of barley cultivar Golden Promise and obtained transgenic lines again the growth conditions were same as described above. Whereas, for thermotolerance assay the plants of transgenic lines were grown at 34 ^◦^C from sowing to three leaf stage and thereafter recovery from heat stress was obtained at 22 ^◦^C, although photoperiod and light intensity was same throughout the experiment.

### Isolation and cloning of *HvHSFA6a* gene

Leaf samples collected at different time points were pooled and RNA isolation was done by using Trizol reagent (Invitrogen, Thermofisher USA). The quality and quantity of isolated RNA was checked on 1% agarose gel and nanodrop (Thermofisher USA), respectively. 2μg RNA and *HvHsfA6a* forward and reverse primers were taken for cDNA synthesis and PCR amplification of 1119 bp ORF of *HvHsfA6a* gene was performed by using one step SuperscriptII RT-PCR system (Invitrogen, Life Technologies, Rockville, MD, USA) according to manufacturer’s instructions. The resulting PCR product was then cloned in pGEMT-easy cloning vector (Promega, USA). Cloning and *HvHsfA6a* gene was confirmed by sequencing followed by BLAST search using NCBI. This gene was further subcloned in *XmaI* forward and *SalI* reverse cloning sites in a blunt cloning PCR vector pJET1.2 (Invitrogen, Thermofisher Scientific USA). Primers used are listed in Supplementary Table 1

### Assay for transcriptional activity and Subcellular localization of HvHSFA6a protein

To check the transactivation potential of HvHsfA6a, yeast system was used. The ORF of *HvHsfA6a* was amplified and fused to GAL4 BD in the yeast vector *pGBKT7* (Clontech, CA, USA), and the resulting constructs was designated as *pGBKT7:HvHsfA6a*. The empty *pGBKT7* was used as a negative control. Plasmids *pGBKT7:HvHsfA6a* was transferred to yeast strain AH109 by lithium acetate-polyethylene glycol-mediated transformation procedure and cultured respectively on the media of SD/−Trp/X-α-Gal (*pGBKT7:HvHsfA6a*) and SD/−Trp (empty *pGBKT7)* at 30 °C for 72–120 h.. Transcriptional activation activity was evaluated by spot and LacZ-filter lift assays.

For subcellular localization, the ORF of *HvHsfA6a* without its stop codon was PCR-amplified by using primer pairs *HvHsfA6a-F-* GGGGACAAGTTTGTACAAAAAAGCAGGCTATGGACGCGGCGGTGCCG and *HvHsfA6a- R*- GGGGACCACTTTGTACAAGAAAGCTGGGTAGAGCGGGCTAGTAGAACTCAGGTACC C and cloned into *pDONR221* and subsequently into binary vector *pB7FWG2* by Gateway^TM^ cloning to construct *pB7FWG2:35S:HvHsfA6a-GFP*. The cloned sequence was verified by sequencing. The preparation of *Agrobacterium* and transient expression in *N. benthamiana* was performed as previously described (Li, 2011). *A. tumefaciens C58* harboring the binary vector *pB7FWG2:35S:HvHsfA6a-GFP* was grown in LB-broth media containing 50 mg/L of spectinomycin, 30 mg/L gentamycin, 30 mg/L of rifampin to the stationary phase at 28 °C. After centrifugation, *A. tumefaciens C58* was resuspended in the infiltration medium (10 mM MgCl_2_, 100 μM acetosyringone) to a final OD_600_ of 0.5. The suspension was infiltrated with a 1-mL needleless syringe into the abaxial surface of 4-week-old leaves of *N. benthamiana*. Protein localization was analyzed 24-48 hours after infiltration by confocal laser scanning microscope (TCS SP8, Leica). GFP was excited at 488 nm and the fluorescence detected between 498-525 nm.

### Construction of vector and barley transformation

Barley HSF *XmaI- SalIHvHsfA6a* was subcloned under ubiquitin promoter (*ZmUbi1*) and *nos* terminator (*NosT*). To achieve this double digestion of empty binary vector *p6oAct-UbiZm-LH* (DNA cloning services) and PCR vector pJET1.2 (Invitrogen, Thermofisher Scientific USA) harbouring gene of interest (*XmaI-SalIHvHsfA6a*) was performed with both the cloning enzymes *XmaI* and *SalI*. Digested products were run on 1% agarose gel. The bands of insert (*XmaI- SalIHvHsfA6a*) and vector (*XmaI-SalI p6oAct-UbiZm-LH*) were excised and extracted using Favorgen Gel/PCR purification kit (Favorgen Biotech. Corp.). The resulting vector and insert were ligated to form a complete cassette *p6oAct-UbiZm-LH:XmaI-SalIHvHsfA6a*. Then the restriction digestion of this vector was performed with *EcoRV* and *PmeI* restriction enzymes to take out the complete cassette containing *ZmUbi1:XmaI-SalIHvHsfA6a:NosT*. This cassette was further cloned in a modified binary vector *pANIC6B* (Mann et al., 2012) and used for *Agrobacterium* (*AGL1*) mediated barley transformation. All the clonings were confirmed by sequencing.

Immature embryos of 14 days after pollination were used as explants for barley transformation. Transformation of immature embryos was performed by vacuum infiltration method described by Hensel et al., (2009) by utilising *Agrobacterium* strain *AGL1* harbouring binary vector *pANIC6B: ZmUbi1:XmaI-SalIHvHsfA6a:NosT*.

### GUS Histochemical, qRT-PCR and immune blot analysis

To confirm the stable transformation event, the intrinsic activity of β-glucuronidase (Gus) was monitored by performing histochemical localization. Tissue samples were taken at different stages i.e. immature embryos, two-week-old calli, six-week-old regenerating calli and plantlet emerging from eight-week-old regenerating calli and leaves of developed plants. These tissue materials were submerged in 2 mL of Gus buffer (0.1 M of sodium phosphate buffer pH7.0, 0.1 M each potassium ferrocyanide and potassium ferricyanide, 10% triton-X-100 and 0.5mg/mL X-gluc. For staining, vacuum was provided for 10 minutes and kept for overnight at 37◦C. For complete removal of chlorophyll samples were destained and stored in 70% ethanol till the photograph taken.

To check the expression of *HvHsfA6a* gene in shoot and root tissues of two-week-old seedlings of barley under control and heat stress conditions and for identification of overexpressing transgenic T0 and T1 lines, total RNA was isolated by using Trizol reagent (Invitrogen, Thermofisher USA) as per manufacturer’s instruction. cDNA was synthesized with 1 μg/μL of RNA concentration using MMLV-RT enzyme (Promega, USA). *HvHsfA6a* gene specific and Actin primers designed by using Primer express 3.0 software (Thermoscientific USA) were used (Supplementary Table 1). Quantitative real time PCR analysis was performed in Quant studio 3 system and by using PowerUp^TM^ SYBR^TM^ Green Master Mix in 96 well plates (Applied Biosystems, Thermofisher scientific, USA). Barley *actin* gene (AK356840.1) was used for normalization and as internal standard. Fold change was calculated by using ^-ddCt^ method.

For protein Western analysis, total leaf protein from WT and transgenic barley were extracted as described previously (Ohnoutkova et al., 2012), with some modifications. Briefly, 200 mg of leaf tissues were grounded in liquid nitrogen and total leaf protein was isolated in extraction buffer (50 mM Tris-HCl; pH 8.0, 0.1% v/v Triton X-100, and 100 mM PMSF) for 1 h at 4 °C. The homogenates were centrifuged at 12,000 rpm for 5 min at 4 °C. The total protein concentrations in supernatant were quantified by the Bradford method (Bradford, 1976) and subjected to immunoblotting. Protein extracts (20 µg) were separated by SDS-polyacrylamide electrophoresis (12% acrylamide) and transferred to PVDF transfer membrane (IPVH00010 Immobilon®-P; MilliporeSigmaTM, Burlington, USA) overnight at 10 mA in a cold room (Mini trans-blot cell; Bio-Rad, Hercules, USA). The membrane was blocked for non-specific sites and incubated with anti-HvHsfA6a in 1:2000 dilutions overnight at 4 °C. After washing three times with TBST, the membrane was incubated with anti-rabbit IgG, HRP linked antibody (7074P2; Cell Signaling Technology, Danvers, USA) in 1:5000 dilutions for 1 h at RT. HvHsfA6a protein was visualized in the membrane by using Pierce™ ECL Western blotting substrate (32109; Thermo Fisher Scientific, Waltham, USA) as per the manufacturer’s protocol.

### Electrolyte leakage index, lipid peroxidation and chlorophyll fluorescence measurement

Relative conductivity-based electrolyte leakage in transgenic lines and wild type plants during heat stress was estimated to assess the membrane stability as described by Hayat et al., (2010). The extent of lipid peroxidation in terms of MDA content was estimated according to the method described by Zheng et al., 2012. Additionally, to assess the effect of heat stress on PSII in same transgenic lines as compared to wild type plants several parameters related to PSII such as maximum photochemical efficiency (*F*_v_/*F*_m_), effective photo chemical efficiency (YII) and photochemical electron transport rate (ETR) were recorded by using Junior-PAM chlorophyll fluorometer (H. Walz, Eifeltrich, Germany) according to Krause and Weis, (1991).

### Staining for ROS, measurement of H_2_O_2_ and enzymatic assays

The leaf sections from two-week old wild type and transgenic barley plants, collected under control and heat stress conditions were stained for superoxide and peroxide species localization following the methodology adapted from Wohlgemuth et al., (2002) and Thordal-Christensen et al., (1997), respectively. For measurement of in situ H_2_O_2_ in leaf tissues, method described by Chakraborty et al., (2016) was followed by using Amplex™ Red Hydrogen Peroxide/Peroxidase Assay Kit (Thermo Fisher Scientific, MA, USA).

For the estimation of anti-oxidative enzyme activities, 500 mg of leaves of three biological replicates of each transgenic and wild type plants were homogenized in 5 mL of one step assay buffer (Phosphate buffer 100 mM pH7.0, Na_2_EDTA 2mM, PVP 1% ) in a pre-chilled mortar and pestle using liquid nitrogen. The homogenate was centrifuged at 22,000 × *g* for 20 minutes at 4 C° .

The supernatant was re-centrifuged at 4 C° . The supernatant (enzyme extract) was collected in fresh 5 mL MCT and stored at ice till the completion of experiment.

SOD: The activity of superoxide dismutase was determined by measuring its ability to inhibit the photochemical reduction of nitroblue tetrazolium chloride, as described by Giannopolitis and Ries, (1977). The assay mixture consisted of 300 μL of the enzyme extract, 100 mM phosphate buffer (pH 7.8), 3 mM EDTA, 200 μM methionine, 2.25 μM nitroblue tetrazolium and 60 μM riboflavin (added at last) in a total volume of 3.2 ml. The tubes were inverted for 2-3 times for proper mixing and placed under fluorescent light. After 15 minutes lights were switched off to stop the reaction. Then the tubes were immediately put in dark. Absorbance was measured at 560 nm, and one unit of SOD activity (U) was defined as the amount of enzyme required to cause 50% inhibition of the nitroblue tetrazolium photoreduction rate. The results were expressed as Umg−1 protein.

Ascorbate peroxidase: The activity of ascorbate peroxidase was assayed according to Nakano and Asada, (1981). The assay mixture consisted of 150 μL of the enzyme extract, 50 mM phosphate buffer (pH 7.0), 0.2 μM EDTA, 0.2 mM ascorbate, and 2.0 mM H_2_O_2_ in a total volume of 3 mL. Ascorbate oxidation was monitored by reading the absorbance at 290 nm at the moment of H_2_O_2_ addition and 1 min later. The difference in absorbance (ΔA290) was divided by the ascorbate molar extinction coefficient (2.8 mM^-^ 1 cm ) and the enzyme activity expressed as μmol of H_2_O_2_ min mg protein, taking into consideration that 1.0 mol of ascorbate is required for the reduction of 1.0 mol of H_2_O_2_ (McKersie and Leshem, 1994).

Catalase: Catalase specific activity was measured according to Aebi et al., (1974). The assay mixture consisted of 100 μL enzyme extract, 100 mM phosphate buffer (pH 7.0) and 200 mM H_2_O_2_ in a total volume of 1.5 mL. The H_2_O_2_ was added in the last and absorbance was measured at 240 nm. The extinction of substrate (H_2_O_2_) by enzymatic decomposition was measured by decrease in absorbance till 3 minutes.

### RNA-Seq and transcriptome analysis

RNA isolation from leaf tissues of five-week-old non-transformed wild type and overexpressing transgenic lines (two replicates of wild type and two transgenic lines) were isolated as per manufacturer’s instruction (Qiagen, Germany). High quality RNA samples with 200 ng/μL concentration were outsourced for RNA-seq. The integrity and purity of extracted mRNA was checked on Bioanalyzer 2100 system (Agilent Technologies, CA, USA). RNA with RNA integrity number (RIN) value above 7.0 was used for construction of cDNA library. Fragmentation of purified mRNA was performed at 90°C in the presence of divalent cations. First strand cDNA synthesis was done using random hexamers and Superscript II Reverse Transcriptase system (Life Technologies, USA). Second strand cDNA synthesis was performed by using first strand as template, DNA polymerase I and RNaseH enzymes. Moreover, the cleaning of cDNAs was done using Beckman Coulter Agencourt Ampure XP SPRI beads.

cDNA libraries for sequencing were constructed using TruSeq RNA Library Prep Kits for Illumina® (NEB, USA) according to the manufacturer’s instructions, followed by indexing of each sample. Clustering of the index-coded samples was performed on a cBot Cluster Generation System using the TruSeq PE Cluster Kit v3-cBot-HS (Illumia) according to the manufacturer’s instructions. Finally, 100 bp paired end sequencing was carried out at Illumina Hi-Seq platform with a read depth of ≥25 p million reads per library.

Quality control of all the reads obtained for each sample was done using NGSQC Tool kit (Patel et al., 2012) and reads with Phred score >Q20 were selected for further analysis. Reference genome of barley from ENSEMBLE (plants/release40/gff3/hordeum_vulgare) was used for alignment of reads and transcript identification. TopHat pipeline (Trapnell et al., 2009) was used for the alignment of reads on reference genome. Cufflink and Cuffdiff pipelines (Trapnell et al., 2009) were used for identification of differentially expressed transcripts in wild type and transgenic replicates using default parameters. Transcripts with log_2_ ratio ≥2 were considered as differentially expressed. Unsupervised hierarchical clustering of differentially expressed genes was done using Cluster 3.0 and visualized using Java Tree View. Gene ontologies and pathways that harbour expressed transcripts were identified using DAVID Functional Annotation Tool (DAVID Bioinformatics Resources 6.8, NIAID/NIH) (Huang et al., 2009).

CuffDiff pipeline was used for analysis of differentially expressed transcripts among samples of wild type and transgenic replicates. The threshold of absolute fold change was set ≥2 and statistically significant P value threshold adjusted for false discovery rate of less than 0.001 by applying Student’s t-test. Statistically significantly enriched functional classes with a P value adjusted for false discovery rate of less than 0.05 derived using the hypergeometric distribution test corresponding to differentially expressed genes were determined using Student’s t-test with Benjamini Hocheberg FDR test. GO ontologies and KEGG pathways analysis was done using DAVID Functional Annotation Tool (DAVID Bioinformatics Resources 6.8, NIAID/NIH) (Huang et al., 2009).

### ChIP-sequencing and analysis

Young leaves from mature WT and transgenic plants overexpression HvHsfA6a were harvested, washed gently with RO water, wiped with soft tissue papers and then chopped in to 2-3 mm pieces. Then the samples were treated with 37% formaldehyde to crosslink the proteins to the DNA. After crosslinking, ChIP (Chromatin Immunoprecipitation) was performed using the reagents and protocol provided in the universal plant ChIP-Seq kit (Cat. No. C01010152; Diagenode, Ougree, Belgium) as per the manufacturer’s instructions. The sheared chromatin was immunoprecipitated using the DiaMag protein A-coated magnetic beads (Diagenode) and 2 µg of anti-HvHsfA6a antibodies. The IP-DNA was given to Ms Bionivid Technology Pvt Ltd (Bengaluru, India) for ChIP sequencing and analysis.

For sequencing sample preparation, the in put and IP DNA were reverse cross-linked with Proteinase K and treated with RNase, followed by DNA isolation using phenol chloroform method. High quality libraries were prepared using NEBNext® Ultra™II DNA Library Prep Kit for Illumina® according to manufacturer’s protocols and paired-end reads of 151bp read length were generated on the Illumina Novoseq 6000 platform.

Foe ChIP Seq data analysis, quality-based filtering and adapter trimming of the raw sequencing reads was done using fastp (v0.20.1). A thresh hold of 30 was set for the phred quality score. This retains only those reads which have 90 bases (60%) with a phred score 30 and above. This ensures that only high-quality data is retained for the analysis. The filtered reads were aligned against the *Hordeum vulgare* (MorexV3_pseudomolecules_assembly) genome from the ensembl plant data base using the Bowtie2 aligner. Non-unique reads (duplicates) were removed from the alignment record using samtools markdup. Peaks were identified using the peak caller Macs2. Peaks were retained if they passed the p-value threshold of 0.05. Annotation resources from the ensembl database were accessed through the R packages biomaRt and ChIPseeker to identify the genes and TSS regions of the peaks. To identify the binding site of the HSF, a motif analysis was performed using HOMER. A region of 1000 bp upstream and 500bp downstream of the peak locations were searched by the findMotifsGenome.pl script in HOMER for potential motif binding sites of length9,11,13 and 15. The genome fasta was provided to the script and it was run with other default settings. HOMER identifies the denovo motifs in the data set and compares against it’s database of known motifs (containing Jaspar resources as well) in the background and identifies enriched motifs.

### Extraction of metabolites

Fresh leaf tissue was collected, washed and air dried at room temperature. Dried leaf tissue (∼500 mg) was crushed in liquid nitrogen and fine powdered samples were used for metabolite extraction following the protocols of Sarkate et al., (2018). One milliliter of pre-cooled extraction mixture [methanol/water/chloroform in the ratio of 2.5:1:1 (v/v/v)] 1 mL) was added to 500 mg powdered samples in a 1.5 mL micro-centrifuge tube followed by vigorous overtaxing for 2 min at room temperature. 4-Phenylphenol (50 µL of from 2 mg/mL methanol stock) was used as internal standard and added into the extraction mixture and then centrifuged at 14000 x g for 5 min. The supernatant (0.8 mL) was collected into a new 1.5 mL micro-centrifuge tube and then 0.4 mL of water was added, followed by 10 S vortexing and then centrifugation at 14000 x g for 5 min. The upper solvent partition phase (methanol/water) was transferred to a new micro-centrifuge tube and then dried out in a vacuum concentrator (Eppendorf Concentrator plus™, Eppendorf; USA) at 200 °C for 2 h followed by 12 h freeze drying in a lyophilizer. Finally, dried material was double derivatized for GC-MS analyses (Lisec et al., 2006). In the first derivatization, 40 μL of methoxyamine hydrochloride (stock solution: 20 mg/ml in pyridine) was added to the dried sample and incubated for 2h at 37 °C. Subsequently, second derivatization was carried out by adding 80 μL of N-methyl-N-(trimethylsilyl)-trifluoroacetamide (MSTFA) and then incubation at 37 °C for 30 min. Derivatization reaction prepared with empty tube without sample served as the control.

### GC-MS analysis

GC- MS analysis was performed on Agilent 7890B gas chromatograph coupled with an Agilent 5977B mass detector (Agilent technologies, CA, USA). 1μL sample was injected into using automatic sampler (7683 B series, Agilent Technologies) in a splitless mode. HP-5 MS column (5% phenyl methyl polysiloxane: 30m x 0.25mm i.d. x 0.25 µm, Agilent technologies) was used for metabolite separation. The temperature program was as follows: Initial temperature of 70 °C for 5 min, was followed by temperature increase to 220 °C at the ramp rate of 10 °C /min and kept on hold for 10 min, finally temperature was increased to 310 °C at the ramp rate of 20 °C /min and kept on hold at 310 °C for 15 min. Total run time calculated was 50 min. Helium was used as carrier gas at a flow rate of 1mL/min. The inlet and MS transfer line temperatures were set to 250 °C and 310 °C respectively. The MS unit was tuned to its maximum sensitivity and the mass range for total ion current was m/z 80-600, and the detector voltage was set at 1700 eV. Each sample was replicated three times. Scan was started after solvent delay of 5 min with scan frequency of 3 S^-1^ (2.0 HZ).

### Metabolite identification

Data acquisition, automatic peak detection, mass spectrum deconvolution, and library search were done by Agilent ChemStation^TM^ software and Wsearch pro (www.wsearch.com.au). The metabolites were identified by matching the mass spectra of target metabolite with the NIST-17 mass spectral library (National Institute of Standards and Technology), and our in-house mass spectral database that include several secondary metabolites, amino acids, organic acids, and sugar standards. Metabolites having matching similarity of 75% or more in library search were only considered. All artifact peaks such as plasticizers, column bleed, derivatizing agent peaks were not considered in final calculation. To obtain accurate peak areas for each metabolite, unique quantification masses were taken and specified. Each mass spectrum was carefully analyzed for co-elution detection.

### Metabolite data pre-processing and statistical analysis

Raw GC-MS data files obtained from Agilent ChemStation^TM^ software were deconvoluted by Automated Mass Spectral Deconvolution and Identification System (AMDIS) using tools available with WsearchPro (www.wsearch.com.au). Resulting data files were converted into .csv (comma separated values) format before uploading in Metaboanalyst 3.0 (http://www.metaboanalyst.ca). Data were normalized using area of internal standard (4- phenylphenol). Data were log transformed with Pareto scaling (mean-centered and divided by the square root of the standard deviation of each variable) before statistical analyses. Statistical significance was calculated by ANOVA (using Fisher’s LSD method; p value < 0.05), We used, PLS-DA analysis (PLS-DA; component 1 vs component 2) to maximize the difference of metabolic profiles between wild type control (WC), wild type heat stress (WH), transgenic control (TC) and transgenic heat stress (TH) samples. The output for PLS-DA data (PC1 vs PC1) were presented as 2D score plot. Variable importance in projection (VIP) scores were used to screen PLS-DA data for important features to distinguish between samples. Heatmaps were created using Metaboanalyst 3.0 by applying Euclidean distance measure, the Ward clustering algorithm, autoscale feature under standardization tool and using normalized data source. A simplified metabolic pathway interaction map was manually created using information from the Kyoto Encyclopedia of Genes and Genomes (KEGG) database via pathway analysis tool in Metaboanalyst 3.0.

## Supplementary data

Supplementary Table S1-Primer list

Supplementary Table S2-list of differential gene expression analysis

Supplementary Table S3-List of motifs and genes identified by ChIP Seq

Supplementary Table S4-motif enrichment by Homer analysis of ChIp seq

Supplementary Table S6 and S7-Fold changes of metabolites

Supplementary Fig. S1-stages in barley transgenic development

Supplementary Fig. S2. qRT-PCR analysis of *HSP* genes

Supplementary Fig. S3 Schematic representation of differentially expressed genes in photosynthetic and respiratory pathways

Supplementary Fig. S4 A schematic representation of differentially expressed genes of jasmonic acid biosynthesis pathway

Supplementary Fig. S5 A schematic representation of differentially expressed genes of flavonoid biosynthesis pathway

Supplementary Fig. S6: Gene Ontology analysis of potential targets of HvHsfA6a identified by ChIP sequencing

Supplementary Fig. S7: Typical GC-MS chromatogram (TIC) of wild type control (WC), wild type heat stress (WH), transgenic control (TC) and transgenic heat stress (TH) barley leaf samples

## Acknowledgement

RC, SB thanks UGC for Junior and Senior research fellowships, PS, SKM and SS thank MHRD for fellowships. PS also thanks the Fonds de recherche du Québec – Nature et technologies (FRQNT) for a research and travel grant. We thank Dr Rahul Kumar and Dr Amanjot Singh for useful discussion.

## Author contribution

HC conceptualized the study; RC, SB, PS, SS, SSS performed experiments; RC, SB, PS, SS, SKM, MSR, HG, DS and HC performed data analysis; all authors wrote and approve the manuscript.

## REFERENCES

1. Aebi, H. (1984). Catalase in vitro. InMethods in enzymology. Academic Press. 105:121–126.

2. Albihlal, W.S., Obomighie, I., Blein, T., Persad, R., Chernukhin, I., Crespi, M., Bechtold, U., Mullineaux, P.M. (2018). Arabidopsis HEAT SHOCK TRANSCRIPTION FACTORA1b regulates multiple developmental genes under benign and stress conditions. 19;69(11):2847–2862 Journal of Experimental Botany.

3. Ali Khan, H. and Mutus, B. (2014). Protein disulfide isomerase a multifunctional protein with multiple physiological roles. Frontiers in Chemistry, 26(**2**):70.

4. Balfagon, D., Sengupta, S., Gómez-Cadenas, A., Fritschi, F.B., Azad, R.K., Mittler, R. and Zandalinas, S.I. (2019). Jasmonic acid is required for plant acclimation to a combination of high light and heat stress. Plant Physiology, 181(**4**):1668–1682.

5. Baniwal, S.K., Bharti, K., Chan, K.Y., Fauth, M., Ganguli, A., Kotak, S., Mishra, S.K., Nover, L., Port, M., Scharf, K.D. and Tripp, J. (2004) Heat stress response in plants: a complex game with chaperones and more than twenty heat stress transcription factors. Journal of Biosciences, 29(**4**):471–87.

6. Banti, V., Mafessoni, F., Loreti, E., Alpi, A. and Perata, P. (2010). The heat-inducible transcription factor HsfA2 enhances anoxia tolerance in Arabidopsis. Plant Physiology, 152(**3**):1471–1483.

7. Bäurle, I. (2016). Plant heat adaptation: priming in response to heat stress. F1000Research, 5.

8. Bita, C. and Gerats, T. (2013). Plant tolerance to high temperature in a changing environment: scientific fundamentals and production of heat stress-tolerant crops. Frontiers in Plant Science, 4:273.

9. Bradford, M.M. (1976). A rapid and sensitive method for the quantitation of microgram quantities of protein utilizing the principle of protein-dye binding. Analytical biochemistry, 72(**1-2**): 248–254.

10. Chan-Schaminet, K.Y., Baniwal, S.K., Bublak, D., Nover, L. and Scharf, K.D. (2009). Specific interaction between tomato HsfA1 and HsfA2 creates hetero-oligomeric superactivator complexes for synergistic activation of heat stress gene expression. Journal of Biological Chemistry. 284(**31**):20848–20857.

11. Charng, Y.Y., Liu, H.C., Liu, N.Y., Hsu, F.C. and Ko, S.S. (2006). Arabidopsis Hsa32, a novel heat shock protein, is essential for acquired thermotolerance during long recovery after acclimation. Plant Physiology, 140(**4**):1297–1305.

12. Chauhan, H., Khurana, N., Agarwal, P. and Khurana, P. (2011). Heat shock factors in rice (*Oryza sativa* L.): genome-wide expression analysis during reproductive development and abiotic stress. Molecular Genetics and Genomics, 286(**2**):171.

13. Chauhan, H., Khurana, N., Agarwal, P., Khurana, J.P. and Khurana, P. (2013). A seed preferential heat shock transcription factor from wheat provides abiotic stress tolerance and yield enhancement in transgenic Arabidopsis under heat stress environment. PLoS One, 8(**11**):79577.

14. Chen, J.H., Chen, S.T., He, N.Y., Wang, Q.L., Zhao, Y., Gao, W. and Guo, F.Q. (2020). Nuclear-encoded synthesis of the D1 subunit of photosystem II increases photosynthetic efficiency and crop yield. Nature Plants, 6(**5**):570–580.

15. Choudhury, F.K., Rivero, R.M., Blumwald, E. and Mittler, R. (2017). Reactive oxygen species, abiotic stress and stress combination. The Plant Journal, 90(**5**):856–867.

16. Cohen, S.P. and Leach, J.E. (2019). Abiotic and biotic stresses induce a core transcriptome response in rice. Scientific reports, 9(**1**):1–11.

17. FAO, IFAD, UNICEF, WFP and WHO. (2018). The state of food security and nutrition in the world 2018. Building climate resilience for food security and nutrition. Rome, FAO.

18. Finka, A., Cuendet, A.F.H., Maathuis, F.J., Saidi, Y. and Goloubinoff, P. (2012). Plasma membrane cyclic nucleotide gated calcium channels control land plant thermal sensing and acquired thermotolerance. The Plant Cell, 24(**8**):3333–3348.

19. Fragkostefanakis, S., Roeth, S., Schleiff, E. and Scharf, K.D. (2015). Prospects of engineering thermotolerance in crops through modulation of heat stress transcription factor and heat shock protein networks. Plant, cell & environment, 38(**9**): 881–1895.

20. Friedrich, T., Oberkofler, V., Trindade, I., Altmann, S., Brzezinka, K., Lämke, J., Gorka, M., Kappel, C., Sokolowska, E., Skirycz, A., Graf, A. (2021). Heteromeric HSFA2/HSFA3 complexes drive transcriptional memory after heat stress in Arabidopsis. Nature Communications, 8;12(1):3426.

21. Giannopolitis, C.N. and Ries, S.K. (1977). Superoxide dismutases: I. Occurrence in higher plants. Plant Physiology, 59(**2**):309–14.

22. Hayat, S., Hasan, S.A., Yusuf, M., Hayat, Q. and Ahmad, A. (2010). Effect of 28- homobrassinolide on photosynthesis, fluorescence and antioxidant system in the presence or absence of salinity and temperature in Vigna radiata. Environmental and Experimental Botany. 69(**2**):105–112.

23. Hensel, G., Kastner, C., Oleszczuk, S., Riechen, J. and Kumlehn, J. (2009). *Agrobacterium*- mediated gene transfer to cereal crop plants: current protocols for barley, wheat, triticale, and maize. International Journal of Plant Genomics, 835608

24. Higashi, Y., Okazaki, Y., Myouga, F., Shinozaki, K. and Saito, K. (2015). Landscape of the lipidome and transcriptome under heat stress in Arabidopsis thaliana. Scientific reports, 5:10533.

25. Hu, J. and Bogorad, L. (1990). Maize chloroplast RNA polymerase: the 180-, 120-, and 38- kilodalton polypeptides are encoded in chloroplast genes. Proceedings of the National Academy of Sciences, 87(**4**):1531-1535.

26. Hu, L., Bi, A., Hu, Z., Amombo, E., Li, H. and Fu, J. (2018). Antioxidant metabolism, photosystem ii, and fatty acid composition of two tall fescue genotypes with different heat tolerance under high temperature stress. Frontiers in Plant Science, 9:1242.

27. Huang, D.W., Sherman, B.T., Zheng, X., Yang, J., Imamichi, T., Stephens, R. and Lempicki, R.A. (2009). Extracting biological meaning from large gene lists with DAVID. Current Protocols in Bioinformatics. 27(**1**):13–1.

28. Huang, Z.H., Wang, Z.L., Shi, B.L., Wei, D., Chen, J.X., Wang, S.L. and Gao, B.J. (2015). Simultaneous determination of salicylic acid, jasmonic acid, methyl salicylate, and methyl jasmonate from Ulmus pumila leaves by GC-MS. International journal of analytical chemistry, 2015.

29. Hwang, S.M., Kim, D.W., Woo, M.S., Jeong, H.S., Son, Y.S., Akhter, S., Choi, G.J. and Bahk, J.D. (2014). Functional characterization of A rabidopsis HsfA6a as a heat-shock transcription factor under high salinity and dehydration conditions. Plant, cell & environment, 37(**5**): 1202–1222.

30. Jagadish, S.V.K., Muthurajan, R., Oane, R., Wheeler, T.R., Heuer, S., Bennett, J. and Craufurd, P.Q. (2010). Physiological and proteomic approaches to address heat tolerance during anthesis in rice (Oryza sativa L.). Journal of experimental botany, 61(1), pp.143–156.

31. Kamauchi, S., Nakatani, H., Nakano, C. and Urade, R. (2005). Gene expression in response to endoplasmic reticulum stress in *Arabidopsis thaliana*. The FEBS Journal, 272(**13**):3461–76.

32. Kaplan, F., Kopka,J., Haskell, D.W., Zhao, W., Schiller, K.C., Gatzke, N., Sung, D.Y., Guy, C.L. (2004). Exploring the temperature stress metabolome of Arabidopsis. Plant Physiology. 1;136(4):4159-68.

33. Katiyar, A., Smita, S., Lenka, S.K., Rajwanshi, R., Chinnusamy, V. and Bansal, K.C. (2012). Genome-wide classification and expression analysis of MYB transcription factor families in rice and Arabidopsis. BMC genomics, 13(**1**):544.

34. Krause, G.H. and Weis, E. (1991). Chlorophyll fluorescence and photosynthesis: the basics. Annual review of plant biology. 42(**1**):313–349.

35. Lämke, J., Brzezinka, K., Altmann, S. and Bäurle, I. (2016). A hit-and-run heat shock factor governs sustained histone methylation and transcriptional stress memory. The EMBO journal, 35(**2**):162–175.

36. Lenzoni, G. and Knight, M.R. (2019). Increases in absolute temperature stimulate free calcium concentration elevations in the chloroplast. Plant and Cell Physiology, 60(**3**):538–548.

37. Li, X. (2011). Infiltration of Nicotiana benthamiana protocol for transient expression via Agrobacterium. Bio-protocol, 1(e95).

38. Li, Z., Zhang, L., Wang, A., Xu, X. and Li, J. (2013). Ectopic overexpression of SlHsfA3, a heat stress transcription factor from tomato, confers increased thermotolerance and salt hypersensitivity in germination in transgenic Arabidopsis. PloS one, 8(**1**):54880.

39. Lin, M.Y., Chai, K.H., Ko, S.S., Kuang, L.Y., Lur, H.S. and Charng, Y.Y. (2014). A positive feedback loop between HEAT SHOCK PROTEIN101 and HEAT STRESS-ASSOCIATED 32- KD PROTEIN modulates long-term acquired thermotolerance illustrating diverse heat stress responses in rice varieties. Plant physiology, 164(**4**):2045–2053.

40. Lisec, J., Schauer, N., Kopka, J., Willmitzer, L. and Fernie, A.R. (2006). Gas chromatography mass spectrometry–based metabolite profiling in plants. Nature Protocols, 1(**1**):387.

41. Liu, H.C. and Charng, Y.Y. (2013). Common and distinct functions of Arabidopsis class A1 and A2 heat shock factors in diverse abiotic stress responses and development. Plant physiology, 163(**1**):276–290.

42. Liu, H.C., Liao, H.T. and Charng, Y.Y. (2011). The role of class A1 heat shock factors (HSFA1s) in response to heat and other stresses in Arabidopsis. Plant, Cell and Environment. 34(**5**):738–51.

43. Liu, J., Yang, H., Lu, Q., Wen, X., Chen, F., Peng, L., Zhang, L. and Lu, C. (2012). PsbP-domain protein1, a nuclear-encoded thylakoid lumenal protein, is essential for photosystem I assembly in Arabidopsis. The Plant Cell, 24(**12**):4992–5006.

44. Liu, X. and Huang, B. (2000). Carbohydrate accumulation in relation to heat stress tolerance in two creeping bentgrass cultivars. Journal of the American Society for Horticultural Science, 125(**4**):442–447.

45. Mann, D.G., LaFayette, P.R., Abercrombie, L.L., King, Z.R., Mazarei, M., Halter, M.C., Poovaiah, C.R., Baxter, H., Shen, H., Dixon, R.A. and Parrott, W.A. (2012). Gateway-compatible vectors for high-throughput gene functional analysis in switchgrass (*Panicum virgatum* L.) and other monocot species. Plant Biotechnology Journal, 10(**2**):226–236.

46. McKersie, B.D. and Leshem, Y.Y. (1994): Stress and stress coping in cultivated plants. Biologia Plantarum, 37(**3**):380–380.

47. Meiri, D. and Breiman, A. (2009). Arabidopsis ROF1 (FKBP62) modulates thermotolerance by interacting with HSP90. 1 and affecting the accumulation of HsfA2-regulated sHSPs. The Plant Journal, 59(**3**):387–399.

48. Mishra, S.K., Poonia, A.K., Chaudhary, R., Baranwal, V.K., Arora, D., Kumar, R., and Chauhan, H. (2020) Genome-wide identification, phylogeny and expression analysis of HSF gene family in barley during abiotic stress response and reproductive development. Plant Gene, 23:100231

49. Mishra, S.K., Tripp, J., Winkelhaus, S., Tschiersch, B., Theres, K., Nover, L. and Scharf, K.D. (2002). In the complex family of heat stress transcription factors, HsfA1 has a unique role as master regulator of thermotolerance in tomato. Genes & Development, 16(**12**):1555-1567.

50. Mittler, R., Vanderauwera, S., Gollery, M. and Van Breusegem, F. (2004). Reactive oxygen gene network of plants. Trends in plant science, 9(**10**):490–498.

51. Mittler, R., Vanderauwera, S., Suzuki, N., Miller, G.A.D., Tognetti, V.B., Vandepoele, K., Gollery, M., Shulaev, V. and Van Breusegem, F. (2011). ROS signaling: the new wave?. Trends in plant science, 16(**6**):300–309.

52. Mizoi, J., Shinozaki, K. and Yamaguchi-Shinozaki, K., 2012. AP2/ERF family transcription factors in plant abiotic stress responses. Biochimica et Biophysica Acta (BBA)-Gene Regulatory Mechanisms, 1819(**2**):86–96.

53. Nakano, Y. and Asada, K. (1981). Hydrogen peroxide is scavenged by ascorbate-specific peroxidase in spinach chloroplasts. Plant and Cell Physiology. 22(**5**):867–80.

54. Narayanan, S., Zoong-Lwe, Z.S., Gandhi, N., Welti, R., Fallen, B., Smith, J.R. and Rustgi, S. (2020). Comparative lipidomic analysis reveals heat stress responses of two soybean genotypes differing in temperature sensitivity. Plants, 9(**4**):457.

55. Nishizawa-Yokoi, A., Yoshida, E., Yabuta, Y. and Shigeoka, S. (2009). Analysis of the regulation of target genes by an Arabidopsis heat shock transcription factor, HsfA2. Bioscience, biotechnology, and biochemistry, 73(**4**):890-895.

56. Nishizawa, A., Yabuta, Y., Yoshida, E., Maruta, T., Yoshimura, K. and Shigeoka, S. (2006). Arabidopsis heat shock transcription factor A2 as a key regulator in response to several types of environmental stress. The Plant Journal, 48(**4**):535–547.

57. Nover, L., Bharti, K., Döring, P., Mishra, S.K., Ganguli, A. and Scharf, K.D. (2001). Arabidopsis and the heat stress transcription factor world: how many heat stress transcription factors do we need?. Cell stress & chaperones, 6(**3**):177.

58. Nover, L., Scharf, K.D., Gagliardi, D., Vergne, P., Czarnecka-Verner, E. and Gurley, W.B. (1996). The Hsf world: classification and properties of plant heat stress transcription factors. Cell stress & chaperones, 1(**4**):215.

59. Ogawa, D., Yamaguchi, K. and Nishiuchi, T. (2007). High-level overexpression of the Arabidopsis HsfA2 gene confers not only increased themotolerance but also salt/osmotic stress tolerance and enhanced callus growth. Journal of experimental botany, 58(**12**):3373–3383.

60. Ohnoutkova, L., Zitka, O., Mrizova, K., Vaskova, J., Galuszka, P., Cernei, N., Smedley, M.A., Harwood, W.A., Adam, V. and Kizek, R. (2012). Electrophoretic and chromatographic evaluation of transgenic barley expressing a bacterial dihydrodipicolinate synthase. Electrophoresis, 33(**15**): 2365–2373.

61. Porter, JR., Moot, D.J. (1998). Research beyond the means: climate variability and plant growth. InCOST77,79,711: International Symposium on Applied Agrometeorology and Agroclimatology: proceedings. Publications Office. 13-

62. Reddy, P.S., Kishor, P.B.K., Seiler, C., Kuhlmann, M., Eschen-Lippold, L., Lee, J., Reddy, M.K. and Sreenivasulu, N. (2014). Unraveling regulation of the small heat shock proteins by the heat shock factor HvHsfB2c in barley: its implications in drought stress response and seed development. PloS one, 9(**3**):89125.

63. Ristic, Z., Momcilovic, I., Bukovnik, U., Prasad, P.V., Fu, J., DeRidder, B.P., Elthon, T.E. and Mladenov, N. (2009). Rubisco activase and wheat productivity under heat-stress conditions. Journal of Experimental Botany. 60(**14**):4003–14.

64. Ruan, Y.L. (2014). Sucrose metabolism: gateway to diverse carbon use and sugar signaling. Annual review of plant biology, 65:33–67.

65. Ruan, Y.L., Jin, Y., Yang, Y.J., Li, G.J. and Boyer, J.S. (2010). Sugar input, metabolism, and signaling mediated by invertase: roles in development, yield potential, and response to drought and heat. Molecular Plant, 3(**6**):942–955.

66. Saidi, Y., Finka, A., Muriset, M., Bromberg, Z., Weiss, Y.G., Maathuis, F.J. and Goloubinoff, P. (2009). The heat shock response in moss plants is regulated by specific calcium-permeable channels in the plasma membrane. The Plant Cell, 21(**9**):2829–2843.

67. Saisho, D. and Takeda, K. (2011). Barley: emergence as a new research material of crop science. 724–727.

68. Salvucci, M.E. and Crafts-Brandner, S.J. (2004). Relationship between the heat tolerance of photosynthesis and the thermal stability of Rubisco activase in plants from contrasting thermal environments. Plant Physiology, 134(4):1460–70.

69. Sarkate, A., Saini, S.S., Teotia, D., Gaid, M., Mir, J.I., Roy, P., Agrawal, P.K. and Sircar, D. (2018). Comparative metabolomics of scab-resistant and susceptible apple cell cultures in response to scab fungus elicitor treatment. Scientific Reports, 8(**1**):1–14.

70. Scharf, K.D., Berberich, T., Ebersberger, I. and Nover, L. (2012). The plant heat stress transcription factor (Hsf) family: structure, function and evolution. Biochimica et Biophysica Acta (BBA)-Gene Regulatory Mechanisms, 1819(**2**):104–19.

71. Schramm, F., Ganguli, A., Kiehlmann, E., Englich, G., Walch, D. and von Koskull-Döring, P. (2006). The heat stress transcription factor HsfA2 serves as a regulatory amplifier of a subset of genes in the heat stress response in Arabidopsis. Plant molecular biology, 60(**5**):759–772.

72. Schult, K., Meierhoff, K., Paradies, S., Töller, T., Wolff, P. and Westhoff, P. (2007). The nuclear-encoded factor HCF173 is involved in the initiation of translation of the psbA mRNA in Arabidopsis thaliana. The Plant Cell, 19(**4**):1329–1346.

73. Serrano, N., Ling, Y., Bahieldin, A. and Mahfouz, M.M. (2019). Thermopriming reprograms metabolic homeostasis to confer heat tolerance. Scientific reports, 9(**1**):1–14.

74. Shen, Z., Ding, M., Sun, J., Deng, S., Zhao, R., Wang, M., Ma, X., Wang, F., Zhang, H., Qian, Z. and Hu, Y. (2013). Overexpression of PeHSF mediates leaf ROS homeostasis in transgenic tobacco lines grown under salt stress conditions. *Plant Cell*, Tissue and Organ Culture (PCTOC), 115(**3**):299–308.

75. Shi, S., Li, S., Asim, M., Mao, J., Xu, D., Ullah, Z., Liu, G., Wang, Q. and Liu, H. (2018). The Arabidopsis calcium-dependent protein kinases (CDPKs) and their roles in plant growth regulation and abiotic stress responses. International journal of molecular sciences, 19(**7**):1900.

76. Shim, D., Hwang, J.U., Lee, J., Lee, S., Choi, Y., An, G., Martinoia, E. and Lee, Y. (2009). Orthologs of the class A4 heat shock transcription factor HsfA4a confer cadmium tolerance in wheat and rice. The Plant Cell, 21(**12**):4031–4043.

77. Song, Y., Chen, Q., Ci, D., Shao, X. and Zhang, D. (2014). Effects of high temperature on photosynthesis and related gene expression in poplar. BMC plant biology, 14(**1**):111.

78. Spreitzer, R.J. and Salvucci, M.E. (2002). Rubisco: structure, regulatory interactions, and possibilities for a better enzyme. Annual Review of Plant Biology. 53.

79. Thirumalaikumar, V.P., Gorka, M., Schulz, K., Masclaux-Daubresse, C., Sampathkumar, A., Skirycz, A., Vierstra, R.D. and Balazadeh, S., 2021. Selective autophagy regulates heat stress memory in Arabidopsis by NBR1-mediated targeting of HSP90. 1 and ROF1. Autophagy, 17(**9**): 2184-2199.

80. Thordal-Christensen, H. and Zhang, Z., Wei, Y., Collinge, D.B. (1997) Subcellular localization of H2O2 in plants. H2O2 accumulation in papillae and hypersensitive response during the barley-powdery mildew interaction. Plant J, **(11)**: 1187-1194.

81. Trapnell, C., Pachter, L. and Salzberg, SL (2009). TopHat: discovering splice junctions with RNA-Seq. Bioinformatics, 25(**9**):1105–11.

82. Vasseur, F., Pantin, F. and Vile, D. (2011). Changes in light intensity reveal a major role for carbon balance in Arabidopsis responses to high temperature. Plant, cell & environment, 34(**9**):1563–1576.

83. von Koskull-Doring, P., Scharf, K.D. and Nover, L. (2007). The diversity of plant heat stress transcription factors. Trends in plant science, 12(**10**):452–457.

84. Wang, X., Xin, C., Cai, J., Zhou, Q., Dai, T., Cao, W. and Jiang, D., 2016. Heat priming induces trans-generational tolerance to high temperature stress in wheat. Frontiers in Plant Science. (7): 501.

85. Wang, W., Jiang, W., Liu, J., Li, Y., Gai, J. and Li, Y. (2017). Genome-wide characterization of the aldehyde dehydrogenase gene superfamily in soybean and its potential role in drought stress response. BMC genomics, 18(**1**):1–17.

86. Wang, X. and Huang, B. (2017). Lipid-and calcium-signaling regulation of HsfA2c-mediated heat tolerance in tall fescue. Environmental and Experimental Botany,136:59–67.

87. Wang, X., Huang, W., Liu, J., Yang, Z. and Huang, B. (2017). Molecular regulation and physiological functions of a novel FaHsfA2c cloned from tall fescue conferring plant tolerance to heat stress. Plant biotechnology journal, 15(**2**):237–248.

88. Wohlgemuth, H., Mittelstrass, K., Kschieschan, S., Bender, J., Weigel, H.J., Overmyer, K., Kangasjärvi, J., Sandermann, H. and Langebartels, C. (2002). Activation of an oxidative burst is a general feature of sensitive plants exposed to the air pollutant ozone. Plant, Cell & Environment, 25(**6**): 717–726.

89. Xue, G.P., Drenth, J. and McIntyre, C.L. (2015). TaHsfA6f is a transcriptional activator that regulates a suite of heat stress protection genes in wheat (*Triticum aestivum* L.) including previously unknown Hsf targets. Journal of Experimental Botany, 66(**3**):1025–1039.

90. Xue, G.P., Sadat, S., Drenth, J. and McIntyre, C.L. (2014). The heat shock factor family from *Triticum aestivum* in response to heat and other major abiotic stresses and their role in regulation of heat shock protein genes. Journal of experimental botany, 65(**2**):539–557.

91. Yang, A., Dai, X. and Zhang, W.H. (2012). A R2R3-type MYB gene, OsMYB2, is involved in salt, cold, and dehydration tolerance in rice. Journal of experimental botany, 63(**7**):2541-2556.

92. Yokotani, N., Ichikawa, T., Kondou, Y., Matsui, M., Hirochika, H., Iwabuchi, M. and Oda, K.(2008). Expression of rice heat stress transcription factor OsHsfA2e enhances tolerance to environmental stresses in transgenic Arabidopsis. Planta, 227(**5**):957–967.

93. Yoshida, T., Ohama, N., Nakajima, J., Kidokoro, S., Mizoi, J., Nakashima, K., Maruyama, K., Kim, J.M., Seki, M., Todaka, D. and Osakabe, Y. (2011). Arabidopsis HsfA1 transcription factors function as the main positive regulators in heat shock-responsive gene expression. Molecular Genetics and Genomics, 286(**5-6**):321–332.

94. Zang, D., Wang, J., Zhang, X., Liu, Z. and Wang, Y., 2019. Arabidopsis heat shock transcription factor HSFA7b positively mediates salt stress tolerance by binding to an E-box-like motif to regulate gene expression. Journal of Experimental Botany, 70(**19**):5355–5374.

95. Zhang, W., Zhou, R.G., Gao, Y.J., Zheng, S.Z., Xu, P., Zhang, S.Q. and Sun, D.Y. (2009). Molecular and genetic evidence for the key role of AtCaM3 in heat-shock signal transduction in Arabidopsis. Plant Physiology, 149(**4**):1773–1784.

96. Zheng, Y.H., Li, X., Li, Y.G., Miao, B.H., Xu, H., Simmons, M. and Yang, X.H. (2012). Contrasting responses of salinity-stressed salt-tolerant and intolerant winter wheat (*Triticum aestivum* L.) cultivars to ozone pollution. Plant Physiology and Biochemistry, (52): 169-178.

